# Single residues in the complexin N-terminus exhibit distinct phenotypes in synaptic vesicle fusion

**DOI:** 10.1101/2024.01.12.575336

**Authors:** Estelle Toulme, Jacqueline Murach, Simon Bärfuss, Jana Kroll, Jörg Malsam, Thorsten Trimbuch, Melissa A. Herman, Thomas H. Söllner, Christian Rosenmund

**Author notes:** **Corresponding authors:** Christian Rosenmund and Thomas H. Söllner. **Conflict of interest statement:** The authors declare no competing interests.

## Abstract

The release of neurotransmitters at central synapses is dependent on a cascade of protein interactions, specific to the presynaptic compartment. Amongst those dedicated molecules the cytosolic complexins play an incompletely defined role as synaptic transmission regulators. Complexins are multidomain SNARE complex binding proteins which confer both inhibitory and stimulatory functions. Using systematic mutagenesis and combining reconstituted *in vitro* membrane fusion assays with electrophysiology in neurons, we deciphered the function of the N-terminus of complexin II (Cpx). The N-terminus (amino acid 1 - 27) starts with a region enriched in hydrophobic amino acids (1-12), which can lead to lipid binding. In contrast to mutants which maintain the hydrophobic character and the stimulatory function of Cpx, non-conservative exchanges largely perturbed spontaneous and evoked exocytosis. Mutants in the downstream region (amino acid 11-18) show differential effects. Cpx-A12W increased spontaneous release without affecting evoked release whereas replacing D15 with amino acids of different shapes or hydrophobic properties (but not charge) not only increased spontaneous release, but also impaired evoked release and surprisingly reduced the size of the readily releasable pool, a novel Cpx function, unanticipated from previous studies. Thus, the exact amino acid composition of the Cpx N-terminus fine tunes the degree of spontaneous and evoked neurotransmitter release.

**Significance Statement:** We describe in this work the importance of the N-terminal domain of the small regulatory cytosolic protein complexin in spontaneous and evoked glutamatergic neurotransmitter release at hippocampal mouse neurons. We show using a combination of biochemical, imaging and electrophysiological techniques that the binding of the proximal region of complexin (amino acids 1-10) to lipids is crucial for spontaneous synaptic vesicular release. Furthermore, we identify a single amino acid at position D15 which is structurally important since it not only is involved in spontaneous release but, when mutated, also decreases drastically the readily releasable pool, a function that was never attributed to complexin.

## Introduction

The transmission of information across synapses in mammalian brains requires tightly controlled neurotransmitter release. Presynaptic proteins orchestrate the fusion of neurotransmitter (NT)-filled synaptic vesicles (SVs) with the plasma membrane to release their contents onto the postsynaptic site. The SNARE (<u>S</u>oluble *<u>N</u>*-ethylmaleimide sensitive factor <u>A</u>ttachment protein <u>RE</u>ceptor) proteins, VAMP2, Syntaxin-1a and SNAP25, form the core SV fusion machinery (Sørensen et al., 2006; Zhou et al., 2015; Zhang et al., 2022). The calcium-sensor synaptotagmin 1, together with another release modulatory protein, complexin (Cpx), clamp or stimulate membrane fusion in absence or presence of Ca^2+^, respectively (DiAntonio and Schwarz, 1994; Fernández-Chacón et al., 2001; Reim et al., 2001; Huntwork and Littleton, 2007; Südhof, 2013; Rizo and Xu, 2015).

Complexins are the best characterized SNARE complex binding proteins (Trimbuch and Rosenmund, 2016). Of the four Cpx isoforms in vertebrates, Cpx I to III are expressed in the brain and the retina, whereas Cpx IV only is found in retinal ribbon synapses (Reim et al., 2005). On one hand, Cpxs inhibit synaptic transmission by arresting spontaneous SNARE complex assembly in autaptic neurons (Xue et al., 2010; Malsam et al., 2020), and on the other hand they facilitate action potential-evoked exocytosis (Reim et al., 2001; Trimbuch et al., 2014). However, it should be noted that discrepancies in the role of Cpx in synaptic transmission at vertebrate synapses have been reported using different experimental approaches (Maximov et al., 2009; Yang et al., 2010, 2013; Kaeser-Woo et al., 2012). Across species, the role of Cpx varies. In *C. elegans* and *Drosophila*, Cpx knock-out (KO) causes an increase in spontaneous release, suggesting a role for inhibiting, or clamping, the fusion of release ready vesicles; but also a decrease in the size of the readily-releasable pool (RRP) and evoked response (Huntwork and Littleton, 2007; Wragg et al., 2013; Cho et al., 2014). In mammalian neurons, Cpx plays a prominently stimulatory role in neurotransmitter release. For example, in neurons where Cpxs I-III are knocked out (Cpx-triple knock out; Cpx-TKO), the evoked response is decreased but neither the spontaneous synaptic release nor the RRP are changed (Xue et al., 2010; Trimbuch et al., 2014). Cross-species rescue experiments indeed suggest that Cpx may have adapted to modulate release through suppression of spontaneous release or by boosting Ca^2+^-triggered release depending on the needs of the synapse (Xue et al., 2009).

Cpx is made up of four main domains: the N-terminal domain (NTD), the accessory helix (AH), the central alpha-helix (CH) and the C-terminal domain (CTD). Each of those domains shows distinct interactions and physiological functions (Xue et al., 2007). The CH binds and stabilizes the partially assembled SNARE complex whereas the AH arrests further SNARE complex assembly (Zhou et al., 2017; Malsam et al., 2020). The CTD of Cpx is involved in lipid interactions, stopping SNARE complex assembly, and has been linked to fusion pore-formation and pore-stabilization (Courtney et al., 2022; Hao et al., 2023). Finally, the NTD is crucial for the facilitatory function of Cpx in mammalian neurons, but its mode of action is still poorly understood (Xue et al., 2007, 2009, 2010). While no structural data are available so far, biochemical and NMR data hint that the NTD may interact with membranes (Zdanowicz et al., 2017) or fold back to interact with the SNARE complex (Xue et al., 2008). In addition, the NTD shows species-specific functions (Xue et al., 2008; Yang et al., 2013). Taken together, further and more precise characterization of Cpx NTD function is warranted.

In this study, we aim to dissect the distinct roles that the NTD of Cpx plays in spontaneous and Ca^2+^-evoked NT release using a structure-function approach. Our results show that targeted mutations of single amino acids lead to distinct functional phenotypes that modulate synaptic neurotransmission and go beyond supporting the stimulatory role of Cpx on Ca^2+^-triggered release.

## Materials and Methods

### Animals maintenance and mouse lines

All mouse experiments were performed in accordance with the regulation of the animal welfare committee of the Charité – Universitätsmedizin Berlin. Time pregnant females were anesthetized and euthanized at E18 according to permission by the Landesamt für Gesundheit und Soziales (LAGeSo) Berlin under the license number G106/20.

### Primary hippocampal cultures

Complexin I-III triple KO neurons were generated as previously described (Xue et al, 2008). Primary murine hippocampal neurons were prepared from E18 embryonic mice of either sex, as described previously (Arancillo et al., 2013). Briefly, hippocampi were dissected, and neurons dissociated by an enzymatic treatment using 25 units per ml of papain for 45 min at 37 °C. For conventional microscopy and western blot experiments, 100×10^3^ neurons/well (35 mm diameter) were plated on WT continental astrocyte feeder layers as high-density culture (Chang et al., 2018). For electrophysiology, low-density cultures of 3×10^3^ neurons/well were seeded on astrocyte micro-islands (35 mm diameter) for autaptic cultures (Arancillo et al., 2013). Astrocyte feeder layers were prepared 1-2 weeks before neuronal seeding, as described previously (Arancillo et al., 2013). After plating, neurons were incubated in Neurobasal-A medium supplemented with 50 µg/ml streptomycin and 50 IU/ml penicillin at 37 °C. For electron microscopy (EM) sapphire disks (6 x 0.12 mm, Wohlwend, art. #1292) were coated with PDL + collagen, and ∼30 x 10^3^ cells/cm^2^ were seeded on an astrocytic feeder layer. Electrophysiological, imaging, western blot or electron microscopy experiments were performed at DIV 14-20.

### Lentiviral constructs and virus production

All lentiviral constructs were generated through the Gibson assembly method (New England Biolabs, Ipswich, MA, USA) with the corresponding cDNAs and with a human synapsin-1 promoter-driven lentiviral shuttle vector (f(syn), based on FUGW (Lois et al., 2002)) that could contain either nuclear-localized (NLS) GFP or RFP that was fused C-terminally to a self-cleaving P2A peptide (Kim et al., 2011) to allow polycistronic translation.

For expression of Complexin II (CpxII) variants within neuronal cells a modified lentiviral vector (Lois et al., 2002) was used in which a human Synapsin-1 promoter, driving the expression of a nuclear GFP and the CpxII variant (WT CpxII, or mutants of CpxII). The cDNAs were coupled via a self-cleaving P2A site leading to a bicistronic expression of the 2 proteins (f(syn)NLS-GFP-P2A-CpxII-WPRE). Lentiviral particles were prepared by the Charité Viral Core Facility as previously described ((Lois et al, 2002); vcf.charite.de). Briefly, HEK293T cells were cotransfected with the shuttle vector f(syn)NLS-GFP-P2A-CpxII-WPRE and helper plasmids, pCMVdR8.9 and pVSV.G with polyethylenimine. Virus containing supernatant was collected after 72 h, filtered, aliquoted, flash-frozen with liquid nitrogen, and stored at -80°C. For infection, about 5×10^5^-1×10^6^ infectious virus units were pipetted onto 1 DIV hippocampal CpxI-III triple KO neurons per 35 mm-diameter well.

### Constructs for biochemical reconstitution assays

For reconstitution into small unilamellar vesicles (SUVs), protein constructs containing full length VAMP2 fused to Glutathione S-Transferase (GST) and Synaptotagmin 1-His_6_ lacking the luminal domain (Syt1, amino acids 57-421) are encoded by plasmids pSK28 (Kedar et al., 2015) and by pLM6 (Mahal et al., 2002), respectively. For reconstitution into giant unilamellar vesicles (GUVs), the t-SNARE complex consisting of Syntaxin1A and SNAP-25B is encoded by pSK306. Synthetic genes encoding full-length rat Syntaxin 1A (1-288) with an N-terminal His_6_-tag followed by a prescission protease cleavage site and full-length mouse SNAP-25 (1-206) with an N-terminal GST-tag followed by a prescission protease cleavage site were subcloned into the bicistronic expression plasmid pETDuet1 resulting in pSK306. The cysteines in the SNAP-25 linker region were replaced by serine residues. Short N-terminal extensions after removal of both tags with prescission protease are highlighted in bold. Syntaxin 1A: ***GPG***MKDRTQELRTAKDSDDDDDVTVTVDRDRFMDEFFEQVEEIRGFIDKIAENVEEV KRKHSAILASPNPDEKTKEELEELMSDIKKTANKVRSKLKSIEQSIEQEEGLNRSSADLRIRKTQHSTLSRKFVEVMSEYNATQSDYRERSKGRIQRQLEITGRTTTSEELEDMLESGNPAIFASGIIMDSSISKQALSEIETRHSEIIKLENSIRELHDMFMDMAMLVESQGEMIDRIEYNVEHAVDYVERAVSDTKKAVKYQSKARRKKIMIIICCVILGIIIASTIGGIFG and SNAP-25: **GPLGCGSSG**MAEDADMRNELEEMQRRADQLADESLESTRRMLQLVEESKDAGIRT LVMLDEQGEQLERIEEGMDQINKDMKEAEKNLTDLGKFSGLSVSPSNKLKSSDAYKKAWGNNQDGVVASQPARVVDEREQMAISGGFIRRVTNDARENEMDENLEQVSGIIGNLRHMALDMGNEIDTQNRQIDRIMEKADSNKTRIDEANQRATKMLGSG.

To generate CpxII point mutants, the QuikChange DNA mutagenesis kit (Qiagen) (Ruiter et al., 2019) and the template encoding wildtype His_6_-CpxII (pMDL80, (Malsam et al., 2020) were used. Thereby, the following CpxII constructs were established: A12W (pJAC20), D15W (pSK258), D15N (pLB37), D15A (pSK255), D15K (pSK259). In addition, native amino acids at the following CpxII positions were replaced by 4-Benzoyl-L-Phenylalanin (BPA *Bachem)* as described by Malsam et al., 2020: M1Bpa (pSK176), F3Bpa (pSK167), V4Bpa (pSK168), M5Bpa (pSK169), A8Bpa (pSK172), L9Bpa (pSK173), G10Bpa (pSK174), G11Bpa (pSK175), A12Bpa (pSK178), T13Bpa (pSK196), K14Bpa (pSK197), D15Bpa (pSK198), M16Bpa (pSK199), G17Bpa (pSK200) and K18Bpa (pSK201). The identity of all constructs was validated by DNA sequencing.

### Protein expression and purification

In general, *Escherichia coli* BL21 (DE3) (Stratagene) cells were transfected with expression vectors encoding the desired protein constructs. At an OD_660_ between 0.6 – 0.8, addition of 0.3 mM isopropyl-β-D-thiogalactopyranosid (IPTG) induced protein expression. Afterwards cells were harvested by centrifugation (3500 x g, 15 minutes, Sorvall H-1200) and lysed using the high-pressure pneumatic processor 110L (Microfluidics). Cell fragments and insoluble material was removed by centrifugation at 60,000 rpm (70Ti rotor, Beckman) for 1 hour and the clarified supernatants were aliquoted and snap-frozen in liquid nitrogen.

Full length VAMP2 (V2) was purified as described previously (Weber et al., 1998; Malsam et al., 2012) with following modifications: Cells were grown in ZYM media (Studier, 2005) and protein expression was induced with 0.3 mM IPTG for 3 hours at 25°C.

Purification of Synaptotagmin-1-His_6_ (Syt1) was performed according to Malsam et al., 2012 with the following modifications: Ni-NTA beads with bound Syt1 were washed three times: 20 ml Mg-ATP wash buffer (25 mM HEPES-KOH, pH 7.5, 400 mM KCl, 10% (w/v) glycerol, 20 mM imidazole, 1% (w/v) Triton X-100, 2 mM Mg-ATP, 3 mM ß-mercaptoethanol (ß-ME)), 20 ml calcium wash buffer (25 mM HEPES-KOH, pH 7.5, 400 mM KCl, 10% (w/v) glycerol, 20 mM imidazole, 20 mM CaCl_2_, 1% (w/v) Triton X-100, 3 mM ß-ME) and 20 ml EGTA wash buffer (25 mM HEPES-KOH, pH 7.5, 100 mM KCl, 10% (w/v) glycerol, 0.5 mM EGTA, 1% octyl-ß-D-glucopyranoside (ß-OG), 3 mM ß-ME). Syt1 was eluted with EGTA wash buffer containing 500 mM imidazole instead of EGTA, diluted twice to lower salt concentration to 50 mM and bound to a MonoS Sepharose column (GE Healthcare). Finally, Syt1 was eluted with a gradient of 50 mM to 1 M KCl in 25 mM HEPES-KOH, pH 7.4, 1 mM Mg-ATP, 100 µM EGTA, 200 mM sucrose, 10% (w/v) glycerol, 1% (w/v) ß-OG and 1 mM DTT.

t-SNARE complexes consisting of full length syntaxin-1A and SNAP25B were co-expressed in BL21-DE3 *E. coli* with 0.5 mM IPTG for 3 hours at 25°C and lysed (50 mM HEPES-KOH, pH 7.4, 400 mM KCl, 10% (w/v) glycerol, 200 mM sucrose). Pre-assembled t-SNARE is bound to GST-beads (lysis buffer, 1% (w/v) Triton X-100, 1.5% (w/v) cholate, 2 mM Mg-ATP, 5mM TCEP), and washed with cleavage buffer (25 mM HEPES-KOH, pH 7.4, 200 mM KCL, 10% glycerol, 200 mM sucrose, 1% (w/v) ß-OG, 5 mM TCEP). After an overnight cleavage with 200U PreScission protease (Cytiva) and addition of fresh GST-beads to bind free GST and PreScission protease, the protein complex is purified by binding to a MonoQ Sepharose column (GE Healthcare) at 90 mM (sample dilution with 25 mM HEPES-KOH, 200 mM sucrose, 5% (w/v) glycerol and 1% (w/v) ß-OG) and eluted with a gradient of 100 mM to 1 M KCl in 20 mM MOPS-KOH (pH 7.4), 200 mM sucrose, 5% (w/v) glycerol and 1% (w/v) ß-OG.

His_6_-CpxII was expressed and purified as described by Malsam et al., 2012, but the expression was performed in BL21-DE3 codon+ *E. coli* for 2 hours at 27°C and no pefablock was used. CpxII Bpa mutants were expressed as described in Malsam et al., 2020 and purified as the wildtype.

The concentrations of purified proteins were determined by SDS-PAGE and Coomassie Blue staining using BSA as a standard and the Fiji software for quantification. For reconstitution assays, mutant and wildtype protein concentrations were directly compared on a single gel.

### Protein reconstitution into liposomes

Atto488-DPPE (1,2-dipalmitoyl-SN-glycero-3-phosphoethanolamine) and Atto550-DPPE were purchased from ATTO-TEC, all other lipids were from Avanti Polar Lipids. For small unilamellar vesicles (SUVs) containing VAMP2 and Syt1, lipid mixes (3 µmol total lipid) with the following composition were prepared: 25 mol% 1-palmitoyl-2-oleoyl-SN-glycero-3-phosphoethanolamine (POPE), 29 mol% 1-palmitoyl-2-oleoyl-SN-glycero-3-phosphocholine (POPC), 25 mol% cholesterol (from ovine wool), 5 mol% PI (L-α-phosphatidylinositol), 15 mol% 1,2-dioleoyl-SN-glycero-3-phosphoserine (DOPS), 0.5 mol% Atto488-DPPE and 0.5 mol% Atto550-DPPE. The t-SNARE GUV lipid mix (5 µmol total lipid) had the following composition: 34 mol% POPC, 15 mol% DOPS, 20 mol% POPE, 25 mol% cholesterol, 4 mol% PI, 1 mol% PI(4,5)P2 (L-α-phosphatidylinositol-4,5-bisphosphate), 0,05 mol% Atto647-DPPE and 0,5 mol% tocopherol.

Synaptotagmin 1 and VAMP-2 were reconstituted into SUVs at a protein to lipid ratio of 1:900 and 1:350 as described previously (Malsam et al., 2020) with following buffer changes. 3 µmol dried lipid mix was resuspended with VAMP2 in 1,7% (w/v) octyl-ß-D-glucopyranoside (ß-OG) containing dilution buffer (25 mM HEPES-KOH, pH 7.4, 550 mM KCl, 100 µM EGTA). After addition of Syt1, small unilamellar Syt1/VAMP2 vesicles were formed by rapid detergent dilution by a three-fold volume increase with ß-OG free dilution buffer. Overnight dialysis (25 mM HEPES-KOH, pH 7.4, 135 mM KCl, 100 µM EGTA, 1 mM DTT) removed the detergent, followed by centrifugation via Nycodenz gradient to concentrate the liposomes. After a second overnight dialysis (25 mM HEPES-KOH, pH 7.4, 135 mM KCl, 10 µM EGTA, 1 mM DTT) removed residues of Nycodenz, SUVs were snap frozen and stored at -80°C.

The reconstitution of t-SNAREs into SUVs and the generation of t-SNARE GUVs by electroswelling have been described previously (Malsam et al., 2012, 2020), respectively) and recent modifications of the protocol are specified in Kádková et al (Kádková et al., 2023). The final lipid to protein ratios were determined by Atto647 fluorescence intensity measurements of the lipids and Coomassie Blue staining of the proteins separated by SDS-PAGE.

### Lipid mixing assay

Membrane fusion (lipid mixing) assays were performed as described previously (Malsam et al., 2012; Kádková et al., 2023). Briefly, t-SNARE-GUVs (14 nmol lipid) were preincubated with VAMP2/Syt1-SUVs (2.5 nmol lipid) in the presence or absence of 6 µM CpxII for 5 min on ice in 100 µL fusion buffer. After transferring samples into a prewarmed 96-well plate (37°C), fluorescence was emitted by measuring Atto488 (._ex_ = 485 nm, ._em_ = 538 nm) at intervals of 10 seconds. Ca^2+^ was added to a final free concentration of 100 µM after 2 minutes. Addition of 0.7% (w/v) SDS and 0.7% (w/v) n-Dodecyl-ß-D-Maltosid (DDM) after 4 minutes stopped the fusion reactions. The measured fusion-dependent fluorescent signals were normalized to the “maximum” fluorescent signal. As a negative control, SUVs were treated with botulinum neurotoxin type D (BoNT-D) to inactivate VAMP2 and the corresponding fluorescence signals were subtracted from individual measurement sets. For each mutant, three independent fusion experiments were performed. To determine the effect of the BPA mutagenesis, the fluorescence signals of the individual mutants were normalized to the wildtype values (Fig. 1A).

**Figure 1:**
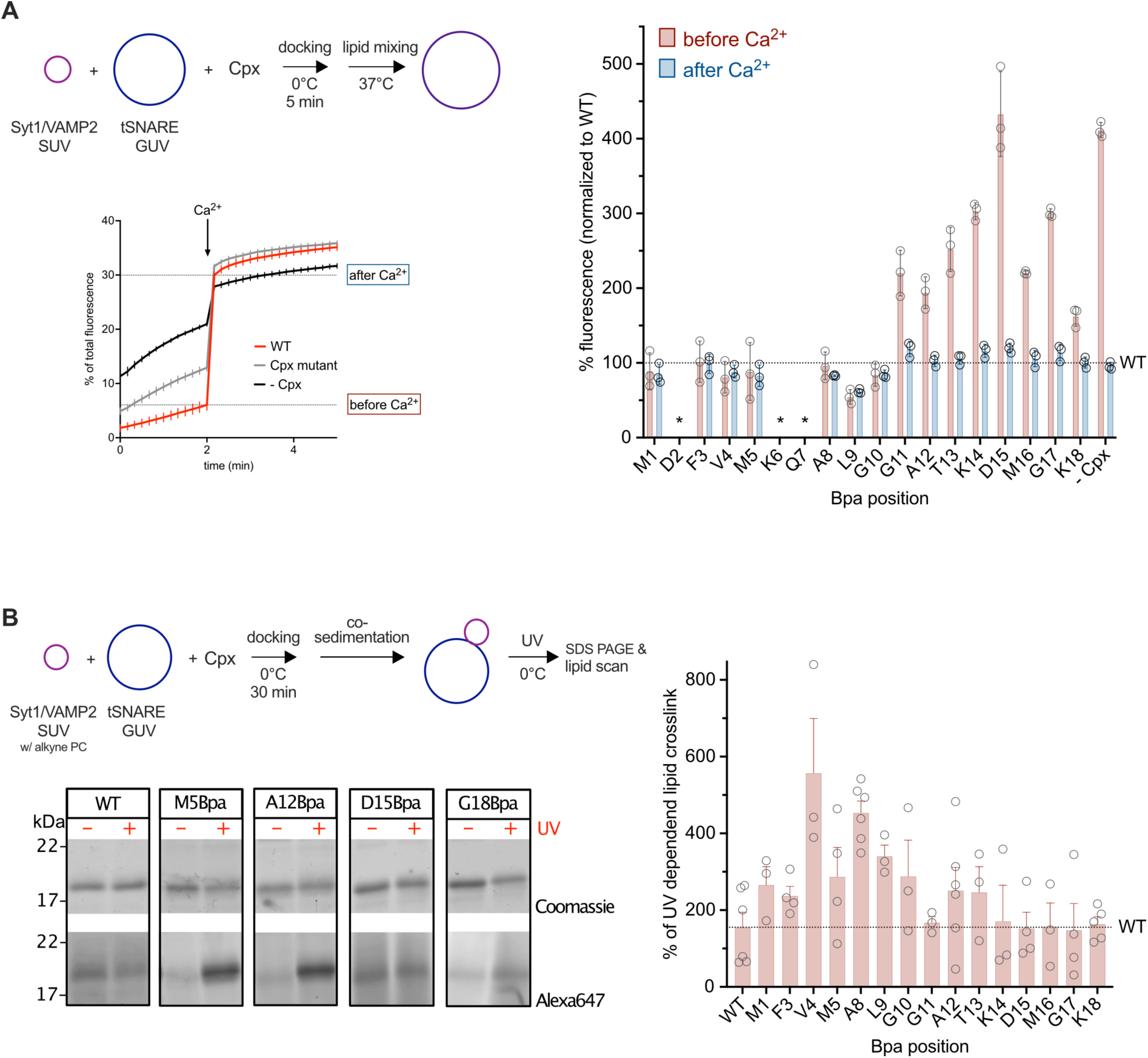
Functional CpxII Bpa mutagenesis screen and lipid crosslinks in the N-terminal domain. **A,** *In vitro* lipid mixing of Syt1/VAMP2 SUVs with t-SNARE GUVs in the absence/presence of Cpx-WT or BPA mutants. Incubation scheme of the reconstituted assay (upper left panel). Representative fusion kinetics show fusion clamping by Cpx (before Ca^2+^ addition) and stimulation after Ca^2+^ addition at 2 min (lower left, the fusion kinetic of CpxII-D15K is shown). Positional Cpx mutants cause distinct effects before (right panel, red bars) and after Ca^2+^ addition (blue bars). All fusion values were normalized to the respective Cpx WT value (100%). * Cpx Bpa variants D2Bpa, K6Bpa, and Q7Bpa could not be purified. Error bars indicate s.e.m with n = 3. **B,** Lipid-crosslinks of positional Cpx Bpa mutants. Experimental incubation scheme is shown in the upper left panel. Representative crosslinks of Cpx mutants to alkyne PC in Syt1/VAMP2 SUVs, as detected by click-labelling of crosslinked alkyne PC with Alexa 647 and SDS PAGE analysis (lower left panel). Quantification of positional lipid crosslinks (right panel). Error bars indicate s.e.m with n = 3-6. Dotted line indicates background lipid crosslinking to Cpx WT, lacking Bpa.

### Detection of lipid interactions via crosslinking of CpxII BPA mutants in an alkyne PC-containing bilayer

For detection of lipid crosslinks (Hempelmann et al., 2023), 22 mol% POPC in the SUV bilayer was replaced by the corresponding amount of alkyne PC (16:0(alkyne)-18:1 PC, Avanti lipids). vSNARE/Syt1-SUVs (5 nmol lipid) and t-SNARE GUVs (28 nmol lipid) were incubated with 6 µM Cpx-WT or BPA mutants in glutathione containing fusion buffer on ice for 30 min to obtain efficient docking and priming as described previously (Malsam et al., 2012). A cushion of 100 µl 70 mM sucrose containing fusion buffer was under layered with 5 µl high sucrose buffer (240 mM sucrose, 1 mM EPPS, 1 mM DTT, pH 8.0) in a separate low-binding tube. After the incubation, samples were layered on top of the 105 µl cushion and spun at 10,000 g for 20 min at 4°C in a swing out rotor (A-8-11 swing bucket rotor, Eppendorf) to isolate CpxII bound to trans SNARE complexes (SNAREpins) formed between the SUVs and GUVs. The supernatant containing free SUVs and soluble CpxII was removed. The pellet was resuspended gently by short vortexing and the suspension was exposed to UV light (365 nm, UV-LED lamp (Opsytec Dr. Gröbel GmbH) with 15 pulses of 1 sec illumination (25 W/cm^2^), pausing for 2 sec between pulses. To fluorescently label the alkyne PC by click chemistry, final concentrations of the following reagents 3125 µM CuSO_4_, 125 µM Tris-[(1-benzyl-1H-1,2,3-triazol-4-yl)-methyl]-amin (TBTA), 3125 µM ascorbic acid and 125 µM Picolyl-Alexa647 (Jena Bioscience) were added to the mix and incubated for two hours at 37°C while smoothly shaking at 400 rpm. Proteins were precipitated by adding 1 mL pre-cooled methanol (-20°C) and incubation overnight at -80°C. After centrifugation (20,000 g, 30 min, 0°C), supernatants were removed, the pellets were dried and resuspended in 12 µl 2x Laemmli buffer. Samples were analyzed by SDS PAGE and scanned for Picolyl-Alexa647 signal before protein staining with Coomassie Blue. Fluorescence intensities corresponding to crosslinked lipids were normalized to respective Coomassie Blue-stained CpxII intensities using Fiji (Fig. 1B).

### High-pressure freezing and freeze substitution

Sample preparation for EM using high-pressure freezing (HPF) and freeze substitution was performed as described previously (Weber-Boyvat et al., 2022). For HPF, neurons were cultured on sapphire glass disks and cryofixed at DIV15-16 using a Leica ICE high-pressure freezer (P > 2,000 bar, cooling rate 10,000-12,000 K/s). After a brief wash in extracellular solution containing 140 mM NaCl, 2.4 mM KCl, 10 mM HEPES, 2 mM CaCl_2_, 4 mM MgCl_2_, and 10 mM glucose (pH adjusted to 7.3 with NaOH, 300 mOsm), sandwiches for HPF were assembled. The sandwiches consisted of the cultured sapphire glass disk, a 100 nm spacer ring, a second blank sapphire glass disk, and a 400 nm spacer ring; the cavity between the sapphire disks was filled with extracellular solution (without cryoprotectants). Directly after the assembly (∼2 min), the samples were high-pressure frozen. The table and chamber of the high-pressure freezer, as well as the extracellular solution were kept at physiological temperature (37°C).

The freeze substitution was carried out using a Leica AFS2 freeze substitution device pre-cooled to -90°C. After the freezing, the sandwiches were disassembled in anhydrous acetone and the samples were subsequently transferred to cryotubes with a freeze substitution solution containing 1 % (w/v) osmium tetroxide, 1 % (v/v) glutaraldehyde, and 1 % ddH_2_O in anhydrous acetone (both steps inside AFS2). The AFS2 was warmed up using the following stepwise heating protocol: -90°C (min. 5 h), -90°C to -20°C (14 h; 5°C/h), -20°C (12 h), -20°C to 20°C (4 h; 10°C/h). At room temperature (RT), the sapphire glass disks were washed 4 x 15 min with anhydrous acetone, stained in 0.1 % (w/v) uranyl acetate in acetone for 1 h, and again washed 4 x 15 min with anhydrous acetone. Epoxy resin (42.8% (w/w) epon, 31.2% (w/w) DDSA, 26 % (w/w) MNA; Fluka) was used for infiltration in ascending concentration: 2 h in 30% epoxy/acetone, 2 h in 70 % epoxy/acetone, and overnight pure epoxy. The following day, the sapphire glass disks were transferred to embedding molds containing epon/araldite (26.3% (w/w) epon, 18.7% (w/w) araldite, 51.9% (w/w) DDSA, 2.98% (w/w) BDMA; EMS) for polymerization (60 °C, 48 h).

### Electron microscopy

Sapphire glass disks were removed from epon blocks using thermal shock. Ultrathin sections (50 nm) were prepared with an Ultracut UCT ultramicrotome (Leica) and collected on formvar-coated 200 mesh copper or nickel grids. The sections were post-stained for contrast enhancement using 2% uranyl acetate (w/v) in ddH_2_O for ∼3 min and Reynold’s lead citrate for ∼1 min. For image acquisition, a FEI Tecnai G2 20 transmission electron microscope (TEM) with an accelerating voltage of 200 kV and equipped with a Veleta 2 x 2K CCD camera (Olympus) was used. Hippocampal synapses were preselected at low magnification to avoid bias and imaged subsequently with a pixel size of 0.71 nm.

### Image analysis

Image analysis and quantification were performed using custom ImageJ macros and python scripts ((Weber-Boyvat et al., 2022); https://github.com/janakroll/synapse-analysis). In ImageJ, electron micrographs of all groups per culture were pooled as image stacks and the order of images was mixed randomly. Only after this randomization, images were excluded if their quality was not good enough for quantification, if the active zone membrane was cut transversally, or if the synapse showed signs of necrosis (swollen or washed out boutons). Approx. 70 images per group and culture were acquired, of which ∼60 images were analyzed.

In ImageJ, the presynaptic active zone (AZ) membrane, docked synaptic vesicles (SVs) and cytosolic SVs were marked. The AZ membrane was defined as the part of the presynaptic membrane that is opposed to the postsynaptic density. SVs were counted as docked SVs if no cleft was visible between outer SV membrane and AZ membrane. Using python, the distribution of SVs was assessed by calculating the shortest distance between SV membranes and AZ membrane.

### Electrophysiology

Whole-cell patch-clamp recordings were performed on autaptic cultures at room temperature at days in vitro 14-21. Synaptic currents were recorded using a Multiclamp 700B amplifier (Axon Instruments) controlled by Clampex 9 software (Molecular Devices). A fast perfusion system (SF-77B; Warner Instruments) continuously perfused the neurons with the extracellular solution contained the following (in mM): 140 NaCl, 2.4 KCl, 10 HEPES (Merck, NJ, USA), 10 glucose (Carl Roth, Karlsruhe, Germany), 2 CaCl_2_ (Sigma-Aldrich, St. Louis, USA), and 4 MgCl_2_ (Carl Roth) (∼300 mOsm; pH 7.4). Somatic whole cell recordings were obtained using borosilicate glass pipettes, with a tip resistance of 2-4 MΩ and filled with the following internal solution (in mM): 136 KCl, 17.8 HEPES, 1 EGTA, 4.6 MgCl_2_, 4 Na_2_ATP, 0.3 Na2GTP, 12 creatine phosphate, and 50 U/ml phosphocreatine kinase (∼300 mOsm; pH7.4). Membrane capacitance and series resistance were compensated by 70% und data filtered by low-pass Bessel filter at 3 kHz and sampled at 10 kHz using an Axon Digidata 1322A digitizer (Molecular Devices).

Neurons were clamped at -70 mV, and action potentials triggered by a 2 ms depolarization to 0 mV to measure EPSCs (excitatory postsynaptic currents). Afterward, EPSCs were induced in the presence of the competitive AMPA receptor antagonist NBQX (3 µM, Tocris Bioscience). Paired-Pulse stimulation for excitatory autapses was assessed by the induction of 2 action potentials with an interstimulus interval of 20 ms. Spontaneous events (mEPSCs) were detected for 40 s. Electrophysiological traces were filtered at 1 kHz, and the range of parameters for inclusion of selected events using a conventionally defined template algorithm in AxoGraph X (AxoGraph Scientific) were 0.15–1.5 ms rise time and 0.5–5 ms half-width. and false positive mEPSC events obtained in NBQX (3 µM) were subtracted to calculate the frequency of spontaneous events. The readily-releasable pool (RRP) was determined by applying an hypertonic 500 mM sucrose solution for 5 s and integrating the transient inward response component (Rosenmund and Stevens, 1996). The Pvr was calculated by dividing the average charge of the EPSC by the RRP charge. Spontaneous release rate was calculated by dividing the mEPSC frequency by the number of synaptic vesicles in the RRP. The number of synaptic vesicles in the RRP was calculated by dividing the RRP size by the mEPSC charge.

In order to estimate synaptic vesicle fusogenicity, we applied on each neuron in paired experiments 500 mM for 5 s or 250 mM sucrose solution for 10 s. The charge transfer of the transient synaptic current was measured and divided by the RRP size from the same neuron to obtain the fraction of RRP released by 250 mM sucrose solution. The onset of sucrose response was identified. We measured the time from the onset of the sucrose solution application to the onset of the response manually (*T1*). We used an “open tip” response configuration to calculate the time needed for the sucrose solution to reach the neuron (*T2*). Subsequently, the sucrose response onset latency was defined as the interval between the time when sucrose solution reaches the neuron and the onset time of sucrose response (*T1* - *T2*). To determine the peak release rate, synaptic current traces were first digitally filtered at 10 Hz. The peak amplitude of sucrose response was then divided by the RRP size from the same neuron to obtain the peak release rate. Data were analyzed offline using AxoGraph X software (AxoGraph Scientific).

### **I**mmunostaining and quantification

Primary hippocampal neuronal mass cultures were rinsed in PBS then fixed for 10 min in 4 % paraformaldehyde (PFA), permeabilized with 0.1 % Phosphate Buffered Saline (PBS)-Tween-20 solution and blocked for an hour with 5 % normal donkey serum. Primary antibodies were used to immune stain overnight at 4°C the following proteins: Complexin (1:1000; Synaptic Systems; #122 002) and VGlut1 (1:4000; Synaptic Systems; #135 304). Subsequently, secondary antibodies labeled with Alexa Fluor 647 anti-guinea pig IgG, Alexa Fluor 488 anti-rabbit and Alexa Fluor 405 anti-chicken raised in donkey serum (each 1:500; Jackson ImmunoResearch) were applied for 1 hour at room temperature, respectively. After washing in PBS, glass coverslips were mounted on glass slides in Mowiol (Polysciences Europe GmbH; Eppelheim, Germany). Images were acquired with an Olympus IX81 epifluorescent microscope equipped with a MicroMax 1300YHS camera using MetaMorph software (Molecular Devices). The analysis was performed offline with ImageJ.

### Western Blot

For detection of Complexin protein levels by western blotting, protein lysates were obtained from mass cultures of Cpx-TKO hippocampal neurons (DIV 14-16) grown on WT astrocyte feeder layers. Briefly, cells were lysed using 50 mM Tris/HCl (pH 7.9), 150 mM NaCl, 5 mM EDTA, 1% Triton-X-100, 1% Nonidet P-40, 1% sodium deoxycholate, and protease inhibitors (complete protease inhibitor cocktail tablet, Roche Diagnostics GmbH; Manheim, Germany). Proteins were separated by SDS-PAGE and transferred overnight at 4°C to nitrocellulose membranes. After blocking with 5% milk powder (Carl Roth GmbH) for 1 hour at room temperature, membranes were incubated with rabbit anti-CpxI/II (1:1000; Synaptic System) and mouse anti-tubulinIII (1:750; Sigma–Aldrich) antibodies for 1 hour at room temperature. The membranes were washed several times with PBS-Tween before being incubated with the corresponding horseradish peroxidase-conjugated goat secondary antibodies (all from Jackson ImmunoResearch Laboratories). Protein expression levels were visualized with ECL Plus Western Blotting Detection Reagents (GE Healthcare Biosciences).

### Structural analysis of complexin mutants

Structures of complexin were visualized in ChimeraX (Pettersen et al., 2021). The sequence of the mutated amino acid was imported from AlphaFold CoLab (Jumper et al., 2021; Mirdita et al., 2022). To generate illustrative representations of Complexin NTD mutants, the NTD was moved into place by hand.

### Experimental design and statistical Analysis

For our electrophysiological and immunocytochemical experiments, we recorded or imaged the same number of neurons per day to minimize variability. Neurons that were analyzed are represented on our bar graph as a single dot. The number of cells recorded or imaged is then explicitly written in the figure legend as well as the number of independent cultures recorded. Statistical tests were performed with Prism 7 (GraphPad software). For bar plots data are represented as mean ± s.e.m. First, all data were tested for normality using the D’Agostino & Pearson test. If they pass the parametric assumption, the Kruskal-Wallis test was performed followed by Dunn’s test. All experiments in the manuscript are repeated at least three times unless explicitly notified. Significance and p values were calculated and reported in the corresponding figure.

### Data and materials availability

Data and Materials of this study are available from the corresponding authors upon reasonable request.

## Results

### Biochemical identification of residues within the N-terminus of Cpx that modify membrane fusion

To assess the role of Cpx’s N-terminal domain (NTD) in regulating spontaneous and Ca^2+^-triggered release we performed a systematic mutagenesis screen, where individual amino acids were mutated and the impact on vesicle fusion was assessed in a lipid mixing assay performed using a reconstituted vesicle fusion assay (Malsam et al., 2012, 2020). We incubated small unilamellar vesicles (SUVs) containing the SV-residing- (v-) SNARE VAMP2, the calcium sensor synaptotagmin-1 (Syt1) and a quenching pair of fluorescence-labeled lipid with giant unilamellar vesicles (GUVs) containing the plasma membrane target- (t-) SNAREs syntaxin-1 and SNAP-25 in the presence or absence of complexin isoform II, CpxII (from here on referred to as Cpx; Fig. 1A, upper left, liposome fusion assay). After a five-minute period of incubation at 0°C, lipid mixing was assessed at 37°C by measuring fluorescence before and after the addition of Ca^2+^. In the presence of wildtype (WT) Cpx, pre-Ca^2+^ fusion, representing spontaneous fusion of reconstituted vesicles, was strongly reduced, observed as an approximately 4-fold reduction in fluorescence signal of lipid mixing (Fig. 1A). The post-Ca^2+^ lipid mixing, reflecting Ca^2+^ triggered fusion, showed a slight tendency to increase in the presence of Cpx WT (Fig. 1A). These results suggest that lipid fusion assays can recapitulate the reported role of Cpx in clamping spontaneous fusion of synaptic vesicles (Xue et al., 2010), but may lack the sensitivity or complex native environment to reflect Cpx’s reported role in facilitating Ca^2+^-triggered release (Xue et al., 2007, 2010).

We next investigated how single residues in Cpx’s NTD affect lipid mixing in reconstituted vesicle fusion assays. To test this, each of the first 18 NTD amino acids of Cpx was exchanged for the photo-activatable, unnatural amino acid benzoyl-phenylalanine (Bpa) (Young et al., 2010; Malsam et al., 2020). Mutants with aspartate at position 2, lysine at position 6, and glutamine at position 7 mutated to Bpa (D2Bpa, K6Bpa, and Q7Bpa, respectively) were insoluble using the heterologous *Escherichia coli* (*E. coli*) expression system and could not be purified. Analogous to a standard mutagenesis approach, we measured the functional contribution of each amino acid with lipid fusion assays *in vitro* without photo-activation before and after the addition of Ca^2+^. With the exception of an inhibition by L9BPA, none of the NTD Cpx mutants affected Ca^2+^-triggered fusion compared to WT. However, Ca^2+^-independent, spontaneous fusion varied by a factor of 4 for the distinct mutants, ranging from 80% to 400% fusion when normalized to WT (Fig. 1A). Generally, the introduction of the bulky and hydrophobic Bpa at amino acids 1-10 had only minor effects on the fusion kinetics. Bpa at positions 11-18 stimulated spontaneous fusion, reaching its largest increase with mutation of the aspartate at position 15 (Fig. 1A right, red bars; before Ca^2+^: 414 +/- 57%, after Ca^2+^: 120 +/- 7%).

While the NTD has been proposed to have an amphipathic structure, we did not observe a specific pattern of activity within the NTD in this assay. Since the NTD of Cpx has been implicated in lipid interactions (Zdanowicz et al., 2017), we investigated which sites are putative lipid interaction sites by using a lipid cross linking assay. Using a setup similar to the lipid mixing assay, the fusion reaction was arrested in the prefusion state and Syt1/VAMP2 SUVs docked to the t-SNARE GUVs were isolated (Fig. 1B). The positional Bpa was photo-activated and crosslinked CpxII–lipid adducts were detected by employing alkyne phosphocholine (alkyne PC) -containing SUV bilayers. To visualize crosslinks, alkyne PC was labeled with a fluorophore (Alexa 647), using click chemistry and the crosslinked products were analyzed by SDS-PAGE and scanning the Alexa 647 fluorescence (Fig. 1B) (Hempelmann et al., 2023). Lower left panel exemplarily shows alkyne PC crosslinks at positions M5Bpa, A12Bpa, D15Bpa by Alexa 647 fluorescence intensities and the corresponding Cpx bands by Coomassie blue staining. Cpx-alkyne PC-Alexa 647 crosslinks are clearly visible for M5Bpa and A12Bpa (Fig. 1B, left panel). For quantification, the fluorescence intensity of Alexa 647 is normalized to the corresponding Cpx amount. Cross-linking between Cpx Bpa mutants and the SUV bilayer was observed throughout most of the N-terminus up to position 13 with prominent crosslinks at positions 4 and 8 (Fig. 1B, right panel), suggesting multiple sites for lipid interactions within the Cpx NTD.

### Cpx-A12W and Cpx-D15W have distinct functional effects on neurotransmitter release

Since our *in vitro* fusion assays suggested a strong role for D15 in regulating spontaneous fusion (Fig. 1A) and a role for A12 in both spontaneous fusion (Fig. 1A) and lipid binding (Fig. 1B), we first investigated these residues for their role in regulating SV fusion in the more complex, native environment of the synapse. For consistency with the biochemical experiments, we mutated these sites to tryptophan, closely mimicking the chemical properties of Bpa substitution (A12W and D15W; Fig. 2B, 2C). In lipid mixing assays, indeed the A12W and D15W mutants exhibited an increase in spontaneous fusion (Cpx-WT: 5.81 +/- 0.62%; A12W: 11.51 +/- 0.57 % and D15W: 18.38 +/- 0.93%; Fig. 3A). Therefore, we went on to perform structure-function experiments on these single point mutants in hippocampal glutamatergic neurons from mice lacking complexin isoforms I, II, and II (Cpx-triple knockout; TKO). Cpx-A12W and -D15W mutants were expressed using lentiviral transduction, and expression levels of re-expressed WT and mutant Cpx were monitored by western blot (WB; Fig. 3.1A) analysis and immunocytochemistry (ICC; Fig. 3.1B). Cpx-A12W and Cpx-D15W proteins were expressed at a similar level as WT-Cpx (Fig. 3-1A), and all three groups were properly localized to presynaptic compartments (Fig. 3-1B).

**Figure 2:**
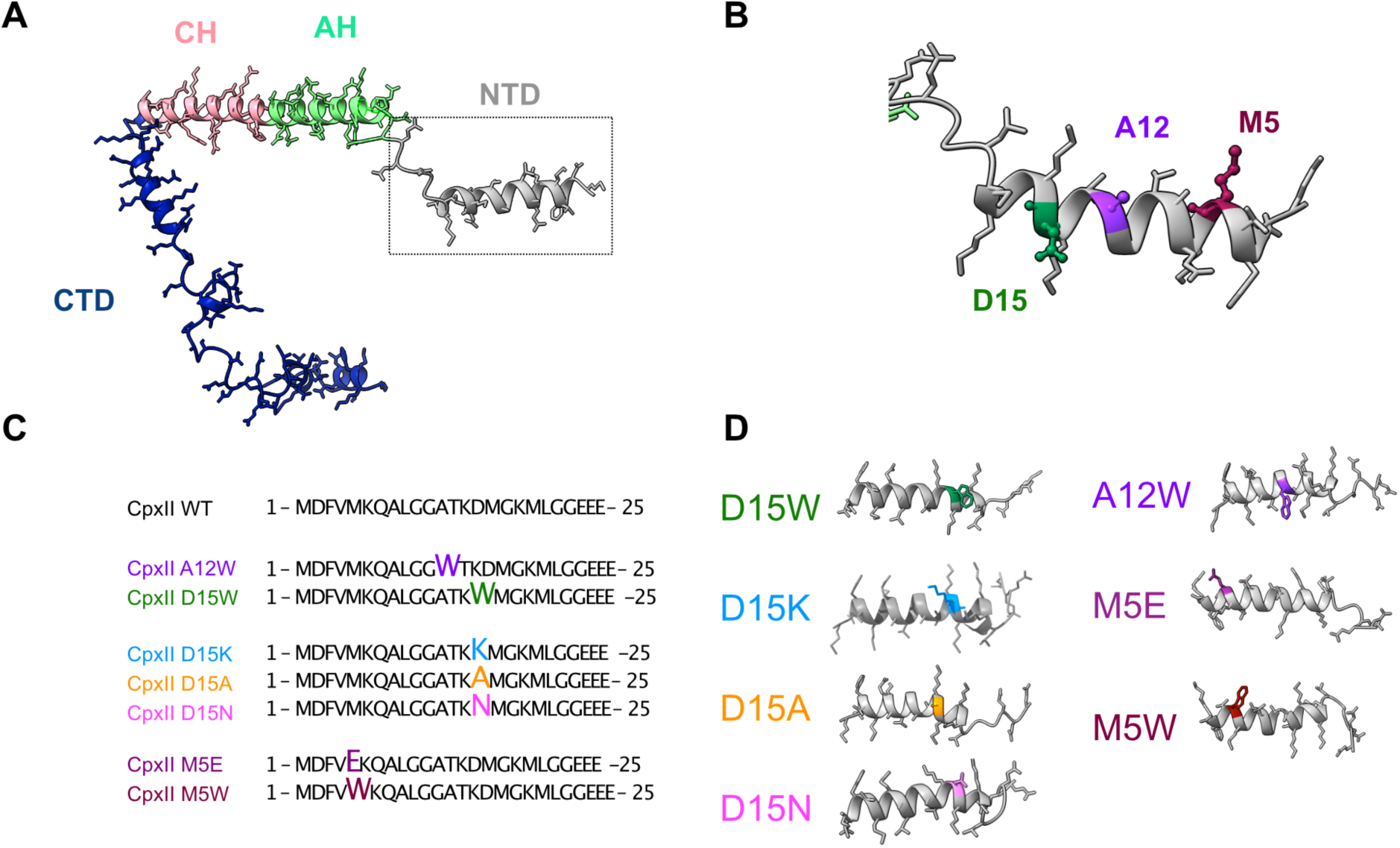
Single mutations of the NTD of CpxII. **A,** Alphafold structure of CpxII whose 4 different domains are color coded, NTD: N-terminal domain (grey), AH: alpha-accessory helix (green), CH: alpha-central helix (light pink) and CTD: C-terminal domain (dark blue). **B,** enlargement of the Alphafold prediction of the structure of CpxII WT NTD highlighting the amino acids of interest for this study. **C,** Sequence alignment of CpxII WT NTD and the various mutants used in this study. **D,** Prediction of the structure of the different mutants of CpxII NTD.

**Figure 3:**
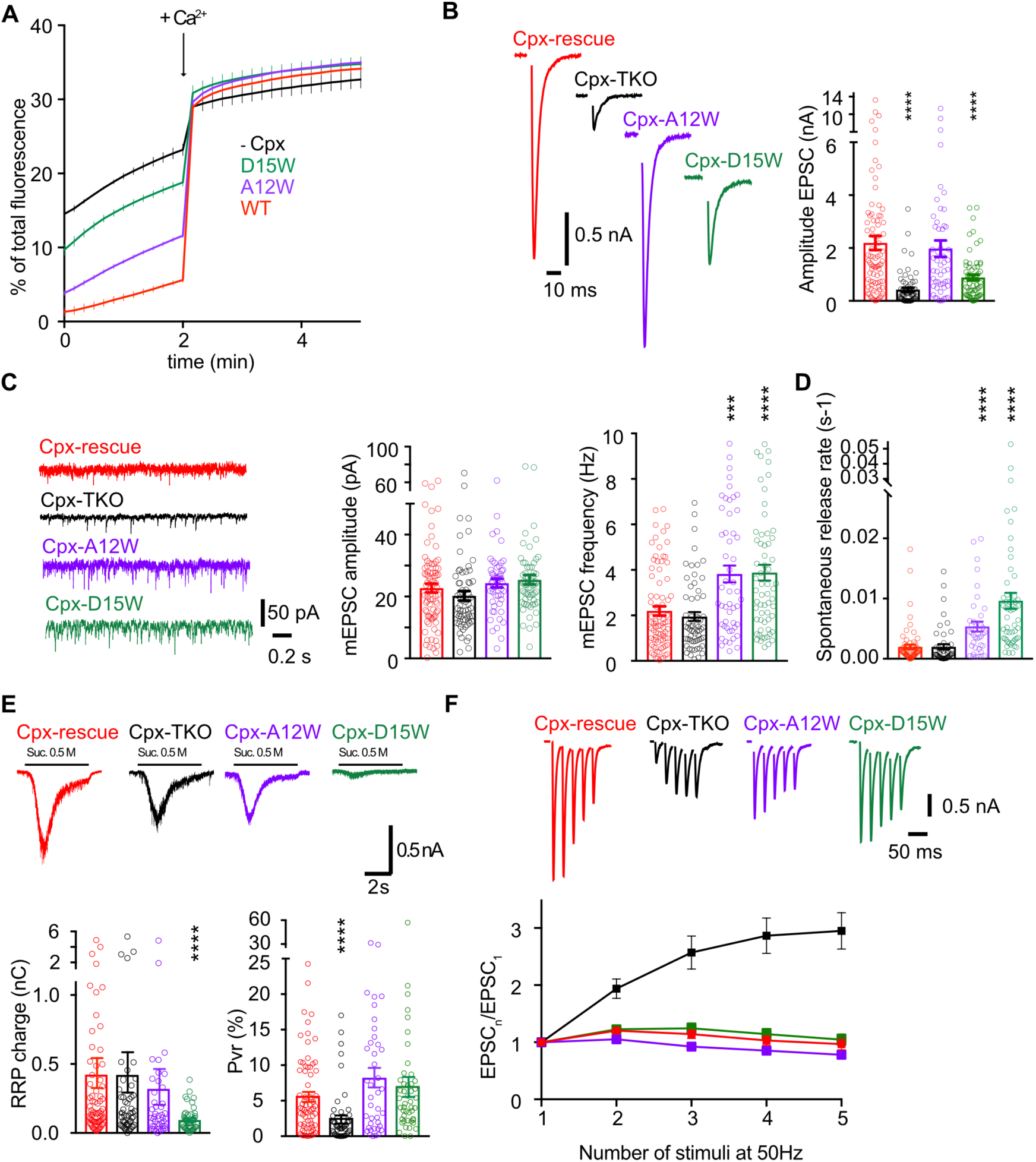
Cpx-A12W and Cpx-D15W have distinct functional effects on neurotransmitter release. **A,** *In vitro* reconstitution assay shows that Cpx-A12W and Cpx-D15W stimulated Ca^2+^-independent lipid-mixing (for experimental set up see Fig. 1A). **B,** Examples traces (left) and quantification of the EPSC amplitude (right) recorded for autaptic hippocampal neurons obtained from Cpx-TKO mice expressing no rescue or rescued with Cpx-WT, Cpx-A12W or Cpx-D15W. **C,** Examples traces (left) and quantification of the miniature EPSC amplitude and frequency (right) obtained from the same neurons as in (B). **D,** Quantification of the spontaneous release rate obtained from the same neurons as in (B). **E,** Examples traces (left) and quantification of the readily releasable pool (RRP) induced by 500 mM sucrose application (right) obtained from the same neurons as in (B). **F,** Examples traces (top) and quantification of short-term plasticity (STP) determined by 50 Hz stimulation and normalized to the first EPSC (bottom) obtained from the same neurons as in (B). The artefacts are blanked in example traces in (B) and (F). The example traces in (C) were filtered at 1 kHz. In (B-E), data points represent a single recorded neuron. Data are expressed as mean ± SEM, asterisks on the graph show the significance comparisons to Cpx-rescue (*** p≤0.001, **** p≤0.0001, nonparametric Kruskal-Wallis test). N = 3 independent cultures.

To further characterize the effects of the A12W and D15W mutations on synaptic transmission, we expressed Cpx-A12W, Cpx-D15W or Cpx WT in Cpx-TKO autaptic mouse culture (Fig. 3B). The phenotypes of each mutant Cpx were compared to the physiological properties of Cpx-TKO and Cpx rescue autaptic neurons (Fig. 3). As previously reported, electrophysiological recordings of cultured, autaptic Cpx-TKO glutamatergic neurons displayed a significantly reduced excitatory postsynaptic current (EPSC) amplitude (Fig. 3B), reduced vesicle released probability (Pvr; Fig. 3E), and facilitation during trains of high frequency stimulation compared to Cpx-WT rescue (Fig. 3; Xue et al., 2010; Trimbuch et al., 2014). Also consistent with previous work, loss of Cpx affected neither the hyperosmotic sucrose evoked pool of fusion competent readily-releasable pool (RRP; Fig. 3E) nor the frequency of miniature EPSCs (mEPSC) (Xue et al., 2010; Trimbuch et al., 2014).

The Cpx-A12W and -D15W mutants showed differential effects on EPSC amplitude rescue as compared to Cpx-WT-rescue (Fig. 3B). Cpx-D15W expressing cells displayed reduced EPSC amplitudes at a level similar to Cpx deficient neurons (D15W: 0.88 +/- 0.1 nA; n = 62; TKO: 0.42 +/- 0.07 nA; n = 67) compared to neurons rescued by Cpx-WT (2.19 +/- 0.26 nA; n = 83), whereas Cpx-A12W rescued to similar levels as Cpx-WT. This suggests that D15 is an important amino acid for the synchronous release of SV. Interestingly, analysis of spontaneous release showed that Cpx-A12W and Cpx-D15W expressing neurons displayed approximately a 2-fold increase in mEPSC frequency compared to Cpx-rescue (Cpx-rescue: 3.89 +/- 0.49 Hz; n = 65; A12W: 7.42 +/- 0.90 Hz; n = 48; D15W: 7.87 +/- 0.67 Hz; n = 48; Fig. 3C) whereas the mEPSC amplitude was similar in all groups (Fig. 3C). The increase in spontaneous release mimics the observed increases in spontaneous lipid mixing seen in the *in vitro* vesicle fusion assay (Fig. 1A and B), and suggests that the A12 and D15 residues in Cpx’s NTD may play a role in clamping spontaneous fusion.

To further examine the role of Cpx’s NTD in spontaneous fusion, we calculated the spontaneous release rate (Fig. 3D), a measure of spontaneous fusion frequency with respect to the number of fusion competent RRP vesicles per cell. Application of hypertonic sucrose revealed that the Cpx-A12W mutant had a similar RRP size compared to Cpx-WT rescue, while Cpx-D15W displayed a drastic reduction in RRP size (A12W:0.321 +/- 0.12 nC; n = 42; D15W: 0.095 +/- 0.012 nC; n = 50; Cpx-rescue: 0.423 +/- 0.100 nC; n = 70; Fig. 3E). The D15W phenotype on RRP size was surprising, as loss of Cpx leads to changes in release efficacy with no effect on RRP size. The spontaneous vesicular release rate, calculated by dividing absolute mEPSC frequency by the number of vesicles in the RRP, showed that spontaneous release was increased for Cpx-A12W and even more robustly for Cpx-D15W (Cpx-rescue: 0.00196 +/- 0.00038 s^-1^; n = 65 versus D15W: 0.00961 +/- 0.00132 s^-1^ n = 54; Fig. 3D), suggesting an unclamping of spontaneous vesicle fusion with both mutations.

To probe whether the observed changes in spontaneous release are related to changes in release efficacy, we calculated synaptic vesicle release probability (Pvr; Fig. 3E) and examined the short-term plasticity (STP) pattern as revealed by a 50 Hz stimulation (Fig. 3F). Pvr, calculated by dividing the charge of the EPSC by the charge of the RRP (Rosenmund and Stevens, 1996), was not significantly different between the Cpx-A12W and Cpx-D15W mutants and Cpx-WT rescue (Cpx-rescue: 5.7 +/- 0.7 %; n = 70; A12W: 8.2 +/- 1.3 %; n = 40; D15W: 7.1 +/- 1.3 %; n = 47; Fig. 3E). Additionally, unlike Cpx-TKO neurons, neither Cpx-A12W nor Cpx-D15W showed a facilitation pattern but rather a depression, similar to Cpx-WT rescue neurons (Fig. 3F). These results demonstrate that the unclamping of spontaneous release related to mutating sites A12 and D15 is not related to altered release efficacy.

### The unique phenotype of Cpx-D15W on spontaneous neurotransmitter release depends on the shape of the amino acid

Because the phenotype of the single point mutation at position D15W of Cpx reduced vesicle priming, revealing a novel phenotype, we investigated whether the decrease in the RRP was due to the loss of the negatively charged amino acid, aspartate (D), or the addition of a hydrophobic amino acid, namely tryptophan (W). To test this, we created three additional single amino acid substitutions for Cpx-D15 (Fig. 2B and 2C) introducing either a neutral and small amino acid (Cpx-D15A), a positively charged amino acid (Cpx-D15K) or polar uncharged amino acid (Cpx-D15N). First, we determined the effects of these novel Cpx-D15 mutants on fusion clamping by analyzing spontaneous and triggered release in the reconstituted liposome fusion assay (Fig. 4A). The mutants affected Ca^2+^-independent membrane fusion in a gradient manner ranging from no clamping (- Cpx) to maximum inhibition Cpx-WT: D15W>D15A>D15K>D15N (-Cpx: 20.68 +/- 0.81%; D15W: 18.31 +/- 1.80%; D15A: 16.09 +/- 1.62%; D15K: 13.32 +/- 1.43%; D15N: 9.91 +/- 2.28% and Cpx-WT: 6.69 +/- 2.00%; Fig 4A). The bulkier or more hydrophobic the amino acid substitution was, the more spontaneous membrane fusion was increased. As the phenotype of Cpx-D15N in *in vitro* experiments is comparable to Cpx-WT, we suggest that the actual charge of the amino acid at position D15 is less critical.

**Figure 4:**
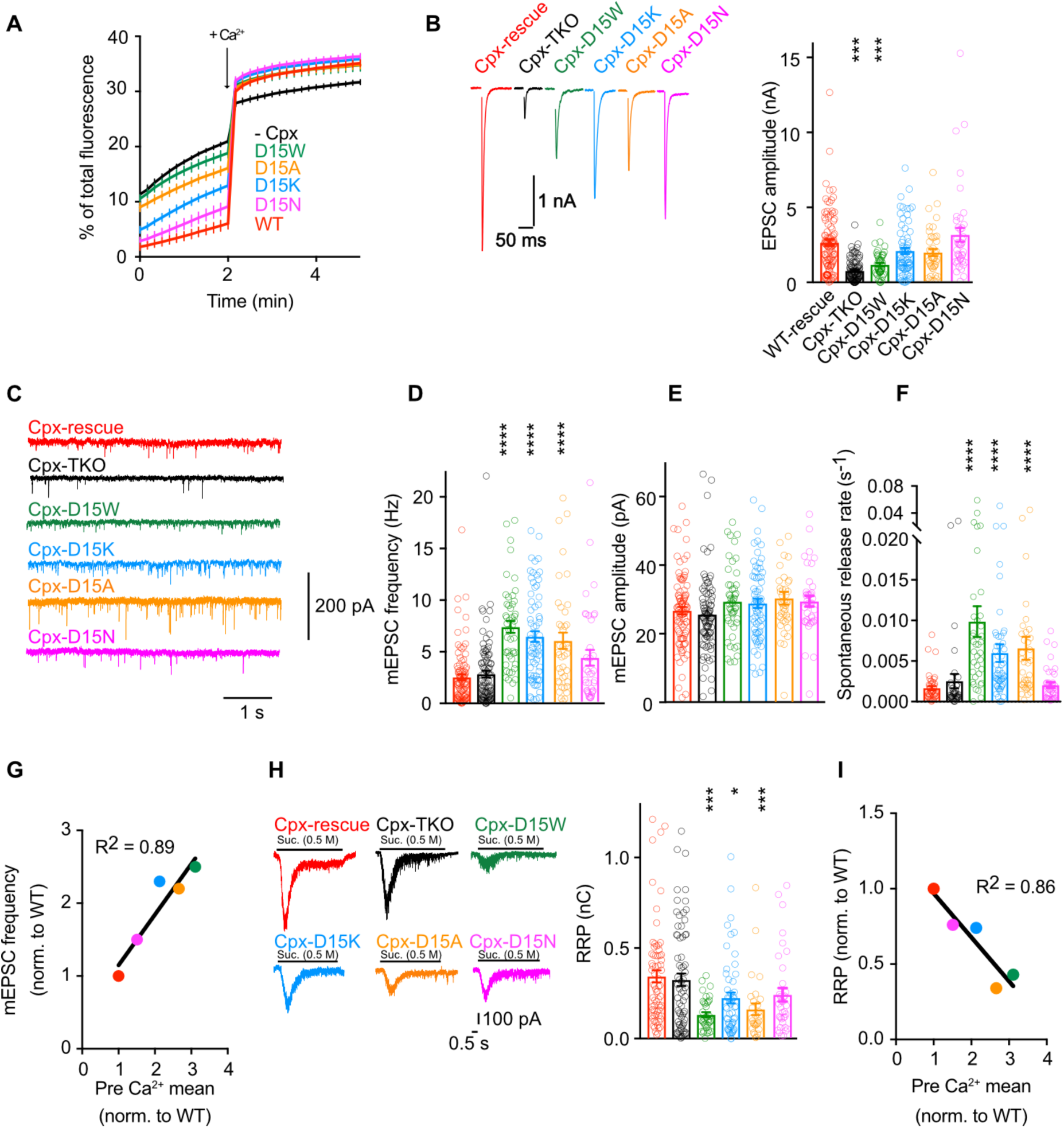
The unique phenotype of Cpx-D15W on spontaneous neurotransmitter release depends on the shape of the amino acid. **A,** *In vitro* reconstitution assay in absence of CPX (- Cpx) or in presence of Cpx-WT or Cpx-D15 mutants with increasing hydrophobicity (D15K > D15N > D15A > D15W), shows lipid mixing kinetics before and after addition of Ca^2+^. Error bars indicate SEM with n = 3 replicates. **B,** Examples traces (left) and quantification of the EPSC amplitude (right) recorded for autaptic hippocampal neurons obtained from Cpx-TKO mice expressing no rescue or rescued with CPX-rescue, Cpx-D15W, Cpx-D15K, Cpx-D15A or Cpx-D15N. **C,** Examples traces of miniature EPSC. **D - F**, Quantification of the mEPSC frequency **(D)**, mEPSC amplitude **(E)** and spontaneous release rate **(F)** obtained from the same neurons as in (B). **G,** Correlative plot of the pre-calcium mean obtained in our biochemical liposome fusion assays against the mEPSC frequency, both normalized to Cpx-rescue values. **H,** Examples traces (left) and quantification of the readily releasable pool (RRP) induced by 500 mM sucrose application (right) obtained from the same neurons as in (B). **I,** Correlative plot of the pre-calcium mean obtained in our biochemical liposome fusion assays against the readily-releasable pool (RRP), both normalized to Cpx-rescue values. The artefacts are blanked in example traces in (B). The example traces in (C) were filtered at 1 kHz. In (B, D, E, F and H), data points represent a single recorded neuron. Data are expressed as mean ± SEM, asterisks on the graph show the significance comparisons to Cpx-rescue (* p≤0.05, ** p≤0.01, *** p≤0.001, **** p≤0.0001, nonparametric Kruskal-Wallis test). N = 3 independent cultures.

We next examined the Cpx-D15 mutants on glutamatergic neurotransmitter release. Expression levels of Cpx-D15 mutants induced by lentivirus in continental hippocampal neuronal cultures assayed by WB (Fig. 4-1A) or ICC (Fig. 4-1B) were comparable to Cpx-WT rescue. In autaptic glutamatergic neurons, evoked EPSCs recorded from Cpx-D15W were decreased to a similar extent as neurons lacking Cpx altogether (Cpx-TKO) when compared to Cpx-WT rescue (Cpx-rescue: 2.66 +/- 0.2 nA; n = 97; Cpx-TKO: 0.77 +/- 0.07 nA; n = 94; D15W: 1.18 +/- 0.11 nA; n = 49; Fig. 4B). However, none of the mutants Cpx-D15K, -D15A or -D15N showed the same decrease in EPSC amplitude (D15K: 2.1 +/- 0.2 nA; n = 69; D15A: 1.93 +/- 0.23 nA; n = 42; and D15N: 3.11 +/- 0.45 nA; n = 42; Fig. 4B). Interestingly, the size of the RRP for Cpx-D15K and Cpx-D15A was still decreased (Cpx-rescue: 0.34 +/- 0.03 nA; n = 70; D15K: 0.22 +/- 0.03; n = 54 and D15A: 0.16 +/- 0.03 nA; n = 30; Fig. 4H) and the mEPSC frequency was still increased (Cpx-rescue: 2.53 +/- 0.28 nA (n = 94), D15K: 6.47 +/- 0.52 (n = 68) and D15A: 5.95 +/- 0.78 nA (n = 41); Fig. 4C – 4F) compared to Cpx Cpx-rescue. Hence, the observed increase in mEPSC frequency or decrease in RRP size was not due only to the loss of a negative charge at position 15 or the inclusion of a hydrophobic amino acid like a tryptophan.

In contrast, mutating D15 into an asparagine (D15N) rescued all the electrophysiological parameters at WT-rescue levels (Fig. 4B-4F and 4H). The RRP size between Cpx-WT rescue and Cpx-D15N were comparable (Cpx-rescue: 0.34 +/- 0.03 nA; n = 70; D15N: 0.24 +/- 0.04; n = 36; Fig. 4H), and the mEPSC frequency in Cpx-D15N neurons was restored to that of the Cpx-WT-rescue (Cpx-rescue: 2.53 +/- 0.28 nA; n = 94, D15N: 4.26 +/- 0.76 nA; n = 38; Fig. 4D). The calculated spontaneous release rate per vesicle was also at Cpx-WT rescue level (Cpx- rescue: 0.0012 +/- 0.0002 s^-1^; n = 78 versus D15N: 0.0019 +/- 0.0003 s^-1^; n = 38; Fig. 4F). These results are in accordance with the *in vitro* experiments performed previously, which showed that Cpx-D15N has an only minor effect compared to Cpx-WT rescue.

To consider these findings altogether we performed correlations between our biochemical experiments and our functional experiments. We observed a strong correlation between the pre-calcium mean release activity obtained through our liposome fusion assay and the mEPSC frequency or the RRP measured in our electrophysiological experiments when these behaviors were plotted against one another for the four D15 mutants and Cpx-WT (mEPSC: R^2^ = 0.89 and RRP: R^2^ = 0.86; Fig. 4G and 4I). This suggests that the D15 residue in Cpx plays a crucial role in clamping spontaneous SV fusion. Additionally, these results strengthen the underlying assumption that pre-calcium fusion rates in the reconstitution assay reflect spontaneous release in mammalian synapses.

### Cpx-D15W partially increases synaptic vesicles fusogenicity

So far, we have shown that the shape of the amino acid at position 15 in Cpx NTD affects spontaneous neurotransmitter release and the size of the pool of SVs ready to be released (RRP) in opposite directions. To further dissect the origin of these regulatory functions, we probed the role of vesicle fusogenicity. The fusogenicity of SVs is a measure of the activation energy required for vesicles to transition from the primed to the fused state, and can be investigated by examining the release kinetics of sucrose-triggered responses in a sucrose-concentration-dependent manner. A 5-second application of 500 mM sucrose represents a saturating stimulus, releasing the entire RRP. In wildtype excitatory neurons an intermediate dose of sucrose (250 mM) releases about a third of the RRP, and changes in the RRP fraction released in 250/500 mM sucrose indicates altered vesicle fusogenicity (Basu et al., 2007; Xue et al., 2010), Fig. 5A). The fraction of RRP released in CPX-TKO neurons is significantly decreased compared to Cpx-WT rescue (consistent with previous observations (Xue et al., 2010); Fig. 5B)). In contrast, Cpx-D15W showed similar vesicle fusogenicity to Cpx-WT rescue (Cpx-rescue: 30.74 +/- 3.57 %; n = 39; TKO: 19.09 +/- 3.67 %; n = 36; D15W: 38.01 +/- 5 %; n = 48; Fig. 5B). To examine vesicle fusogenicity by release kinetic analysis, we quantified the response onset latency of the 250 mM sucrose stimuli, which is slowed down when fusogenicity is lowered. On one hand, the onset response latency for Cpx-D15W was slightly faster compared to Cpx-WT rescue (Cpx-rescue: 0.64 +/- 0.05 s; n = 39 versus D15W: 0.5 +/- 0.04 s; n = 48; p = 0.033; Fig. 5D) consistent with an increase in fusogenicity. On the other hand, the peak release rate which corresponds to the amplitude of the peak response divided by the RRP size is not significantly changed between Cpx-D15W and Cpx-WT rescue (Cpx-rescue: 0.64 +/- 0.07 s^-1^; n = 39; D15W: 0.42 +/- 0.05 s^-1^; n = 48; Fig. 5E).

**Figure 5:**
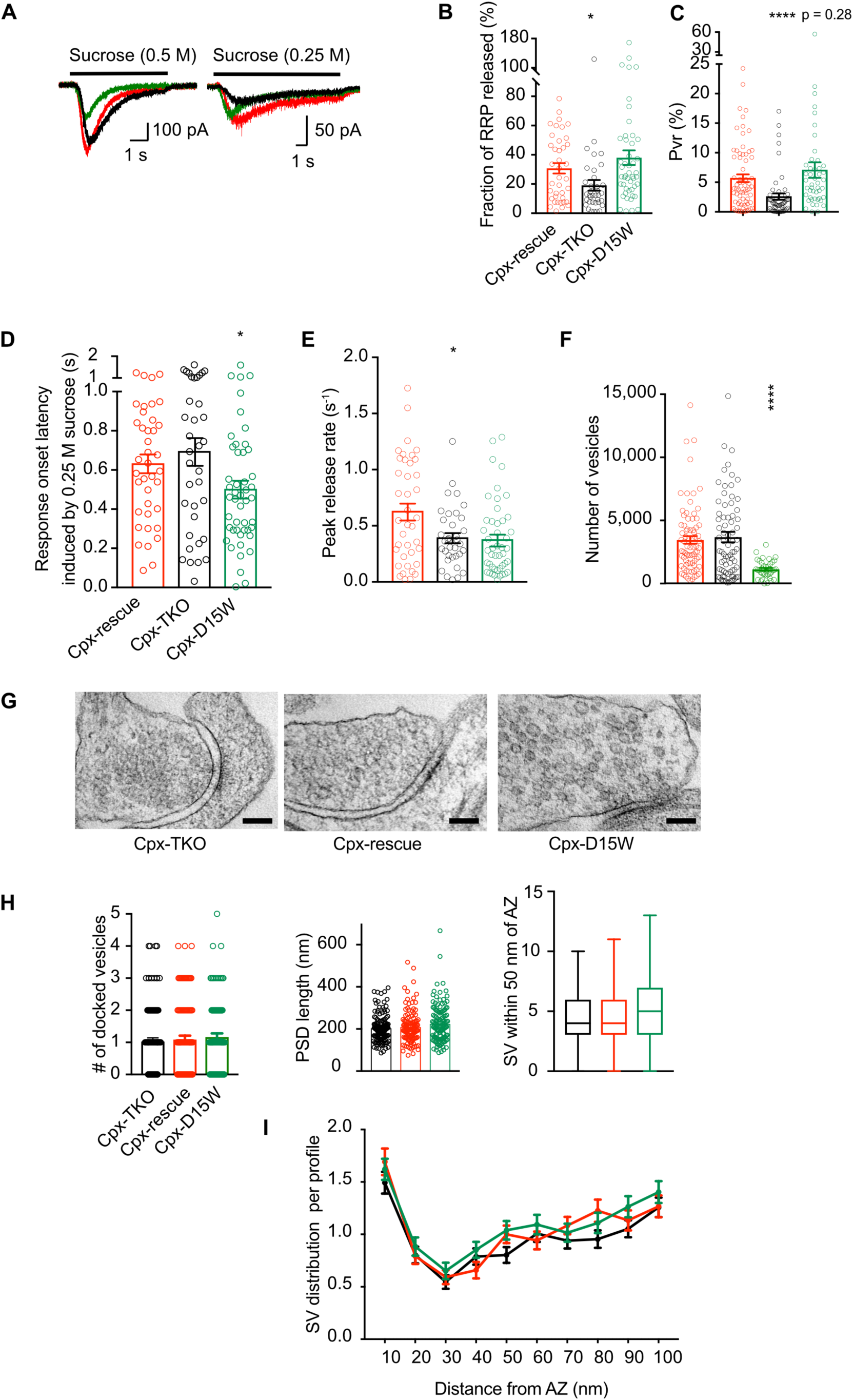
Cpx-D15W partially increases synaptic vesicles fusogenicity. **A,** Example traces of the readily releasable pool obtained from the application of 500 mM and 250 mM sucrose application from Cpx-TKO, Cpx-rescue and Cpx-D15W neurons. **B,** Quantification of the fraction of RRP released determined as the ratio from the charge transfer for 500 mM sucrose and the charge transfer for 250 mM sucrose obtained from Cpx-TKO, Cpx-rescue and Cpx- D15W neurons. **C,** Quantification of probability of vesicular release (Pvr) determined as the percentage of the RRP released upon one AP obtained from the same neurons as in (B). **D,** quantification of the response onset latency determined from the application of 250 mM sucrose and obtained from the same neurons as in (B). **E,** Quantification of the peak release rate determined from the application of 250 mM sucrose and obtained from the same neurons as in (B). **F,** Quantification of the number of vesicles released obtained by dividing the RRP charge measured by 500 mM sucrose application by the mEPSC charge for the same neuron. **G,** Example of high-pressure freezing fixation followed by electron microscopy (HPF-EM) images of neurons from high-density cultures of Cpx-TKO, Cpx-rescue and Cpx-D15W neurons. Scale bar, 200 nm. **H,** Quantification of number of docked vesicles, PSD length, number of synaptic vesicles (SV) within 50 nM of active zone (AZ) obtained for Cpx-TKO, CPX-rescue and Cpx-D15W. **I,** SV distribution within 100 nm of AZ obtained from the same neurons as in (H). In (B-F), data points represent a single recorded neuron. Data are expressed as mean ± SEM, asterisks on the graph show the significance comparisons to Cpx-rescue (* p≤0.05, **** p≤0.0001, nonparametric Kruskal-Wallis test). N = 3 independent cultures.

We observed that the Cpx-D15W mutant has an impaired RRP, as shown by a decreased charge evoked by 500 mM sucrose, compared to either Cpx-WT rescue or Cpx-TKO (Fig. 3E). To translate the impairment of SV priming to the number of fusion competent SVs we divided the RRP charge by the mean charge of mEPSC response (Fig. 3C and 3E), and confirmed that Cpx-D15W expressing neurons have a 3-fold smaller number of RRP SVs compared to Cpx-WT rescue or Cpx-TKO neurons (Cpx-rescue: 3455 +/- 302; n = 78; TKO: 3682 +/- 414; n = 78 and D15W: 1130 +/- 114; n = 38; Fig. 5F). To investigate whether the vesicle priming defect in the D15W mutant translates to an ultrastructural defect, we obtained electron micrographs from synapses and analyzed vesicle docking (Watanabe et al., 2013; Imig et al., 2014). Neurons were plated at high density and fixed using high-pressure freezing followed by freeze substitution, epon embedding and sectioning to 50 nm thickness. Electron microscopic transmission images were analyzed as shown previously (Weber-Boyvat et al., 2022) and the number of docked SVs, considered as those in direct contact with the plasma membrane, were quantified as a function of active zone length. All three groups showed normal postsynaptic density length (Fig. 5G - H). Moreover, the density of docked vesicles was not different between Cpx-WT rescue, Cpx- TKO and Cpx-D15W groups (Cpx-rescue: 1.1 +/- 0.1; n = 120, TKO: 1.05 +/- 0.1; n = 132 and D15W: 1.19 +/- 0.05; n = 129; Fig. 5H). Similarly, the SV distribution profiles within 100 nm of the active zone were comparable between Cpx-WT rescue, Cpx-TKO and Cpx-D15W. These results suggest that the general synaptic organization of neurons lacking Cpx or neurons expressing Cpx-D15W mutant is unaltered. The discrepancy between the low number of vesicles measured with electrophysiology and the absence of a docking phenotype measured via electron microscopy suggests that Cpx-D15W impairs SV priming downstream of SV docking.

### Cpx-D15W unclamps spontaneous vesicular release in WT neurons

While vesicle priming in Cpx-D15W mutant is impaired, vesicle fusogenicity and spontaneous release is enhanced in comparison to Cpx-WT rescue (Fig. 3 and 4). To test whether these effects are dominant, we overexpressed Cpx-WT and Cpx-D15W mutants using lentiviral transduction in wildtype neurons. Overexpression of Cpx-WT in WT neuronal cultures had no measurable impact on any of the electrophysiological parameters we studied in comparison with WT neurons expressing GFP (Fig. 6). Cpx-D15W overexpression, also had no effect on evoked EPSC responses (GFP: 3.51 +/- 0.57 nA; n = 35; Cpx-WT: 3.66 +/- 0.49 nA; n = 35; Cpx-D15W: 4.12 +/- 0.62 nA; n = 37; Fig. 6A). However, the frequency of mEPSC events increased only when Cpx-D15W was overexpressed (GFP: 5.14 +/- 1.18 Hz; n = 34; Cpx-WT: 5.28 +/- 1 Hz; n = 34; and Cpx-D15W: 7.95 +/- 1.19 Hz; n = 34; Fig. 6D) suggesting that Cpx- D15W has a dominant positive effect on spontaneous synaptic transmission. Surprisingly, neither the RRP size, the SV release probably, nor the short-term plasticity dynamics during a 10 Hz action potential train were modified by an overexpression of Cpx-D15W (Fig. 6B, C, E). Taken together the analysis shows that the unusual D15W mutant phenotype is only dominant in unclamping spontaneous release.

**Figure 6:**
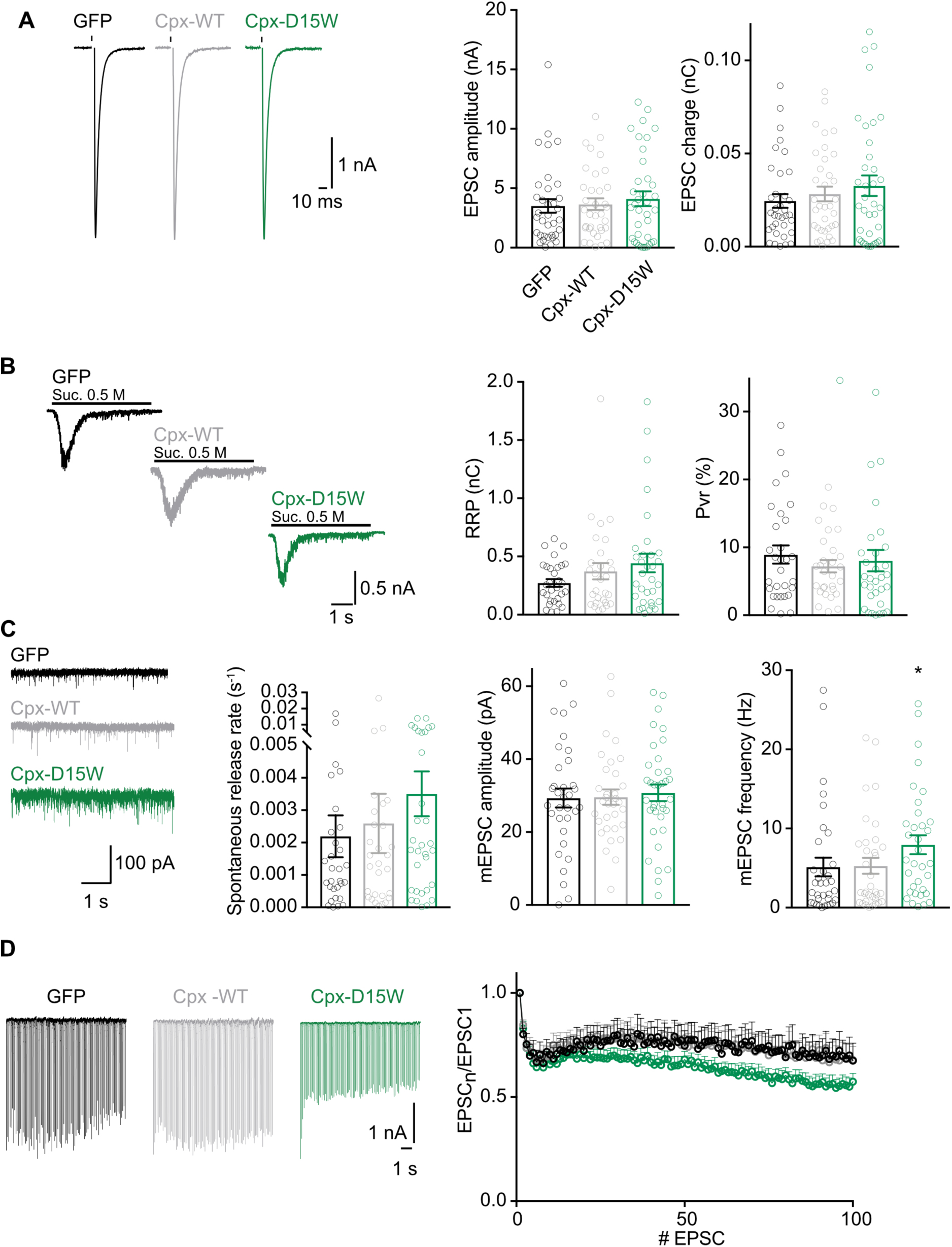
Cpx-D15W has a dominant positive effect on spontaneous vesicular release in WT neurons. **A,** Examples traces (left) and quantification of the EPSC amplitude or EPSC charge (right) recorded for autaptic hippocampal neurons obtained from Cpx-WT mice overexpressing GFP, Cpx-WT or Cpx-D15W. **B,** Examples traces (left) and quantification of the readily releasable pool (RRP) induced by 500 mM sucrose application and probability of vesicular release (Pvr) (right) obtained from the same neurons as in (A). **C,** Examples traces (left) and quantification of the spontaneous release rate, miniature EPSC amplitude and frequency (right) obtained from the same neurons as in (A). **D,** Examples traces (left) and quantification of short-term plasticity (STP) determined by high-frequency stimulation at 10 Hz and normalized to the first EPSC (right) obtained from the same neurons as in (A). The artefacts are blanked in example traces in (A) and (D). The example traces in (C) were filtered at 1 kHz. In (A-C), data points represent a single recorded neuron. Data are expressed as mean ± SEM, asterisks on the graph show the significance comparisons to Cpx-rescue (* p≤0.05, nonparametric Kruskal-Wallis test). N = 3 independent cultures.

### Cpx-M5E mutant disrupts the binding of Cpx N-terminus to the SNARE complex in a charge-dependent manner

Our initial screen with lipid fusion assays predicted the effects of Cpx mutants on regulating spontaneous SV fusion, but suggested no effect of Cpx on Ca^2+^-triggered fusion. This is in contrast to many studies demonstrating the facilitatory effect of Cpx on Ca^2+^ triggered SV fusion in the synapse (Reim et al., 2001; Trimbuch et al., 2014). For instance, our previous work reported that mutating the N-terminal residues Cpx-M5E,K6E disrupts the positive effect of Cpx on SV fusogenicity and synchronous release (Xue et al, 2010). This phenotype was accompanied by a disruption of binding of Cpx NTD to the SNARE complex in nuclear resonance measurements (Xue et al., 2010). Our liposome fusion assay did not show a phenotype in the Cpx-M5-Bpa mutant (Fig. 1A). As the mutation of M5 into a negatively charged glutamic acid was crucial to disrupt the interaction of complexin with the C-terminus of SNARE complex (Xue et al., 2010), we utilized Cpx-M5E mutants in parallel with Cpx-M5W that sterically best mimic the insertion of the unnatural amino acid Bpa.

The Cpx-M5E mutant (Fig. 2B and 2C) was sufficiently expressed and colocalized with the presynaptic glutamatergic marker VGlut1 (Fig. 7-1A and 7-1B). In our functional experiments, Cpx-M5E largely mimics neurons lacking Cpx (Cpx-TKO). The amplitude of the evoked EPSC response is significantly decreased compared to Cpx-WT rescue (Cpx-rescue: 5.37 +/- 0.57 nA; n = 48; TKO: 0.87 +/- 0.12 nA; n = 47 and M5E: 1.49 +/- 0.2 nA; n = 49; Fig. 7A). Moreover, Cpx-M5E showed paired-pulse facilitation comparable to Cpx-TKO (Cpx-rescue: 1.06 +/- 0.13; n = 46; TKO: 1.59 +/- 0.12; n = 46 and M5E: 1.65 +/- 0.11; n = 46; Fig. 7A) and an impairment of SV spontaneous release as shown by the mEPSC frequency decrease compared to Cpx-WT rescue and Cpx-TKO (Cpx-rescue: 3.4 +/- 0.6 Hz; n = 41; TKO: 3.09 +/- 0.44 Hz; n = 42; and M5E: 1.9 +/- 0.8 nA; n = 27; Fig. 7B). On the other hand, the RRP of Cpx-M5E was not different from the RRP for Cpx-WT rescue or Cpx-TKO neurons. In contrast to the phenotype of the Cpx-M5E mutant, when we mutated M5 into a tryptophan (Cpx-M5W; Fig. 2B and 2C), all measured electrophysiological parameters (Pvr, RRP, mEPSC amplitude and STP) behaved like wildtype (Fig. 7A-D). These results are in full accordance with our biochemical experiments, in which Cpx-M5Bpa had no effect on fusion kinetics (Fig. 1A). Taken together, these results support the previously published hypothesis that Cpx NTD residue M5 is crucial for Cpx facilitatory function (Xue et al., 2010), but additionally suggest that the unmasking of this role depends on the charge of the mutated amino acid.

**Figure 7:**
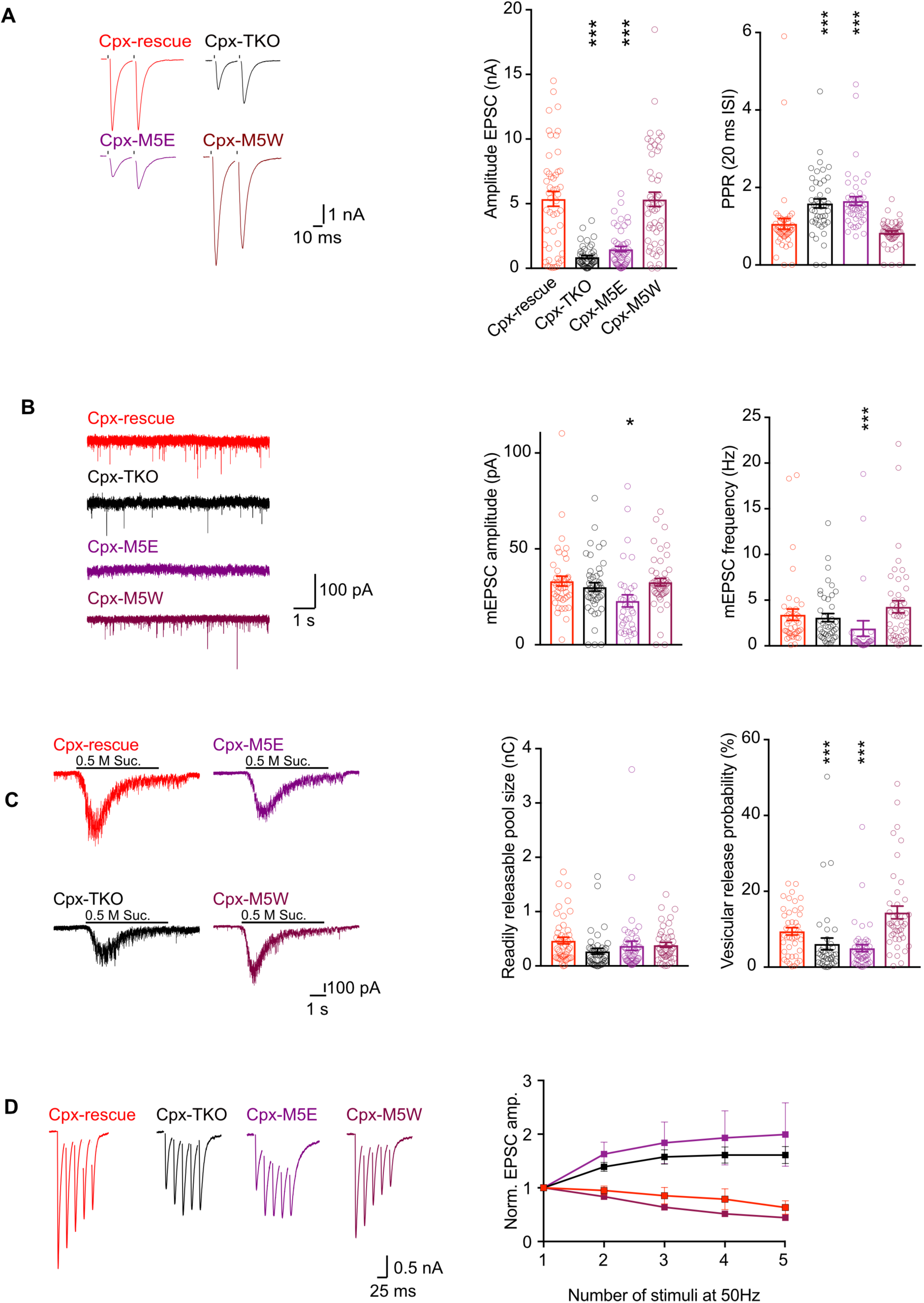
Insertion of a negative charge at position 5 (Cpx-M5E) disrupts the stimulatory function of the Cpx N-terminus. **A,** Examples traces (left) and quantification of the EPSC amplitude (right) and paired-pulse ratio (PPR) at 20 ms intervals recorded for autaptic hippocampal neurons obtained from Cpx-TKO mice expressing no rescue or rescued with Cpx-WT, Cpx-M5E or Cpx-M5W. **B,** Examples traces (left) and quantification of the miniature EPSC amplitude and frequency (right) obtained from the same neurons as in (A). **C,** Examples traces (left) and quantification of the readily releasable pool (RRP) induced by 500 mM sucrose application (right) obtained from the same neurons as in (A). **D,** Examples traces (left) and quantification of short-term plasticity (STP) determined by 50 Hz stimulation and normalized to the first EPSC (bottom) obtained from the same neurons as in (A). The artefacts are blanked in example traces in (A) and (D). The example traces in (B) were filtered at 1 kHz. In (A-C), data points represent a single recorded neuron. Data are expressed as mean ± SEM, asterisks on the graph show the significance comparisons to Cpx-rescue (*p≤0.005, *** p≤0.001, nonparametric Kruskal-Wallis test). N = 3 independent cultures.

## Discussion

The SNARE complex is at the core of the canonical release machinery common for all regulated release. At the synapse, additional modulators are needed to accelerate and tune release properties to adjust the release function to the need of synaptic communication. Complexin is the best characterized SNARE complex binding protein (Ishizuka et al., 1995; McMahon et al., 1995; Takahashi et al., 1995). It can perform both release-accelerating as well as clamping functions, depending on species and synapse (Trimbuch and Rosenmund, 2016 for review). All structural and mutagenesis studies show that its accelerating and clamping functions are embedded in the N- and C-terminal domains (Xue et al., 2009, 2010; Buhl et al., 2013). The NTD is so far the only region known to be essential for the accelerating function of Cpx (Xue et al., 2009, 2010), but how the NTD regulates release is poorly understood. In this study we combined Cpx mutagenesis with *in vitro* reconstitution assays and functional analysis at central synapses to define functionally critical regions within the NTD as well as the putative role of lipid binding of the assumed amphipathic helical secondary structure. We show that i) the proximal region of the N-terminus of Cpx from aa 1 until aa 12 is a lipid interacting region (Fig. 1), ii) residues between 11-16 are important for reducing spontaneous release, iii) mutating the aspartate at position 15 results in a drastically decreased readily releasable pool (Fig. 3 – 6), iv) position M5 is functionally relevant in promoting NT release in synapses (Fig. 7) and sensitive to lipid interaction (Fig. 1). Overall, our detailed analysis of the NTD of Cpx shows that Cpx function is even more complex than initially thought, as the facilitatory NTD can also modulate vesicle priming and spontaneous release.

Overall, our data add significant knowledge to our understanding of Cpx’s mechanisms of action, especially on its facilitatory role. Based on previous studies, the N-terminus, formed by the residues 1-27, is essential for Cpx’s positive action on vesicle release probability (Xue et al., 2010). From previous work in the field, two possible interaction modes for the N-terminus emerge; binding to the SNARE complex itself or interacting with lipid membranes. NMR experiments using a paramagnetic probe at position 12 demonstrated that the N-terminus can interact with the C-terminus of all three SNARE complex forming proteins (Xue et al., 2010; Choi et al., 2018). Assuming an alpha-helical structure of the N-terminus, the helix would contain an amphipathic surface that could also favor membrane binding, where the hydrophobic site is formed by evolutionary conserved residues 1, 5, 8, 9, 12 and 16. Our systematic assessment of lipid interactions using single residue Bpa mediated crosslinking (Fig. 1) showed that the strongest interactions with lipids take place between residues 1-12, but the assay was not selective for residues of the hydrophobic site. Our *in vitro* fusion assays with single residue Bpa mutant Cpx did not display any signs of activation of fusion for the first ten residues (Fig. 1). Thus, changing already hydrophobic residues in this region to Bpa does not affect Cpx function. In contrast, single Bpa mutations at D2, K6, Q7 rendered Cpx insoluble in the heterologous *E. coli* expression system, indicating that the position of hydrophobic amino acids is critical. However, this is likely due to the property of the fusion assay, as deletion of the entire NTD only impairs Ca^2+^-triggered release in mammalian synapses, but not in *in vitro* fusion assays (Bera et al., 2022). Moreover, our Cpx structure-function analysis at the synapse could only identify methionine at position 5 as essential for the facilitatory role of Cpx, suggesting that this residue may carry the role of Cpx’s NTD in increasing synaptic vesicle release probability.

The *in vitro* fusion assay revealed a novel role of Cpx’s N-terminus in fusion clamping, as mutating residues 11 to 17 showed an enhanced pre-calcium fusion activity (Fig. 1). We could confirm that this behavior in pre-calcium fusion activity ascertained in our biochemical analysis indeed reflects a clamping function of spontaneous release, as residues 12 and 15 when mutated and expressed in Cpx-TKO neurons also showed an increase in spontaneous synaptic vesicle fusion (increase in mEPSC frequency) (Figs 3 and 4). Specifically, we observed that spontaneous release was enhanced when residues 12 and 15 were made bulky and hydrophobic and the capability to form hydrogen bonds was impaired. Another important observation linking the behavior of proteins driving fusion within the biochemical assays and within the native habitat of the synapse was the strong correlation between pre-calcium fusion activity in lipid mixing assays and the degree of spontaneous release recorded with electrophysiology in mutants with graded effects on the phenotype (Fig. 4G). These results strongly suggest the fundamental nature of Cpx’s role in SNARE-mediated membrane fusion, as the phenotype is reflected in both in a situation containing the minimal players involved in the process (reconstituted liposome fusion assays) and in an environment containing far more players than necessary for fusion (native synapse). In turn, taken together these results emphasize the impact of the Cpx-NTD on clamping spontaneous release.

One striking observation we made through our detailed analysis of the Cpx-NTD was the novel, single-residue dependent role of Cpx in determining the pool of primed, fusion competent vesicles. We found that altering the acidic nature of the aspartic acid at position 15 resulted in a strong reduction in the RRP (Figs 3 and 4). The effect was not accompanied by a defect in SV docking (Fig. 5), a phenotype typically associated with priming deficiency, arguing that the impairment in vesicle priming occurs at a step downstream of vesicle docking. Comparison of our functional experiments to our data from the *in vitro* assay also showed that the impact on mEPSC frequency and RRP were inversely correlated (Fig. 4G and 4I). Such a behavior points towards a destabilized primed vesicle state, where primed vesicles transition more easily into the unprimed or fused state. Interestingly, we observed a similar behavior in syntaxin 1 and SNAP25 mutants (Salazar-Lázaro et al., 2023) that form the “primary interface” of interaction between synaptotagmin1 and the SNARE complex as defined by protein crystallography (Zhou et al., 2015). In a recent publication, Hao et al show using optical tweezers that removal of the NTD from Cpx changes the disassembly between Cpx and the SNARE complex (Hao et al., 2023), emphasizing the role of the NTD in stabilizing the SNARE complex.

In summary, our analysis of the Cpx N-terminus showed a diverse and complex regulation of release function where single residues mutations in close proximity have discrete and distinct effects on vesicle priming, spontaneous release and the efficacy of Ca^2+^-triggered release. A possible explanation for such behavior is that Cpx, along with synaptotagmin, the SNARE complex and the membranes interact at a closely confined space. The spatial position of the NTD in the tripartite interface, for example in relation to the SNARE complex and synaptotagmin, still needs to be structurally determined. However, our study contributes to a better understanding of the role of the small cytosolic regulator, Cpx, in the fine-tuning of vesicle release, and ultimately in regulation of synaptic transmission.

## Acknowledgements

This work was funded by the Deutsche Forschungsgemeinschaft (DFG; German Research Foundation) project 278001972-TRR186 (to E.T., Ja.M., S.B., T.T., T.H.S., and C.R.). We are grateful to Berit Söhl-Kielczynski, Bettina Brokowski, Katja Pötschke, Ursula Göbel, Lara Braun, Susanne Kreye and Heike Lerch for excellent technical assistance. We also thank Denisa Jamecna and Doris Höglinger for sharing their expertise in lipid crosslinking using click chemistry and Daniela Schweinfurth and Britta Brügger for sharing their knowledge in protein-lipid extraction. We thank the services of the Charité viral core facility for virus production and the electron microscopy core facility for technical support. Molecular graphics were performed with UCSF ChimeraX, developed by the Resource for Biocomputing, Visualization, and Informatics at the University of California, San Francisco, with support from the National Institutes of Health R01-GM129325 and the Office of Cyber Infrastructure and Computational Biology, National Institute of Allergy and Infectious Diseases.

## Author contributions

E.T. performed electrophysiological, biochemical and immunocytochemical experiments in neurons

Ja.M., Jo.M. and S.B., performed *in vitro* biochemical experiments

J. K. performed electron microscopy experiments

E.T., Ja.M., T.H.S. and C.R. designed experiments

T.T. designed lentiviral constructs

E.T., Ja.M., M.A.H., T.H.S. and C.R. wrote and edited the manuscript

---

**Figure 3-1: Protein and synaptic expression of Cpx-A12W and Cpx-D15W mutants in mice hippocampal neuron cultures. A,** Example image (left) of a Western Blot against complexin (top) and Tubulin (bottom) for Cpx-WT rescue and Cpx mutants in Cpx-TKO hippocampal neuron cultures. Bar graph (right) corresponds to the intensity of the complexin bands in each condition normalized to WT (n = 3 independent cultures). **B,** Example images (left) and quantification (right) of immunofluorescence labeling for Complexin and V-Glut1 for continental hippocampal cultures of Cpx-TKO neurons or infected with Cpx-rescue, Cpx-A12W or Cpx-D15W. Scale bar, 10 μm. The quantification of the immunofluorescence intensity of Complexin is normalized to the immunofluorescence intensity of V-Glut1. The values are normalized to the one obtained for Cpx-WT rescue (red). In (B), data points represent a field of view. Data are expressed as mean ± SEM, asterisks on the graph show the significance comparisons to Cpx-rescue (*** p≤0.001, nonparametric Kruskal-Wallis test). N = 3 independent cultures.

**Figure 4-1: Protein and synaptic expression of Cpx-D15 mutants in mice hippocampal neuron cultures. A,** Example image (left) of a Western Blot against complexin (top) and Tubulin (bottom) for Cpx-WT rescue and Cpx mutants in Cpx-TKO hippocampal neuron cultures. Bar graph (right) corresponds to the intensity of the complexin bands in each condition normalized to WT (n = 4 - 5 independent cultures). **B,** Example images (left) and quantification (right) of immunofluorescence labeling for Complexin (green) and V-Glut1 (red) for continental hippocampal cultures of Cpx-TKO neurons or infected with CPX-rescue, Cpx-D15K, Cpx-D15N, Cpx-D15A or Cpx-D15W. Scale bar, 10 μm. The quantification of the immunofluorescence intensity of Complexin is normalized to the immunofluorescence intensity of V-Glut1. The values are normalized to the one obtained for Cpx-WT rescue. In (B), data points represent a field of view. Data are expressed as mean ± SEM, asterisks on the graph show the significance comparisons to Cpx-rescue (**** p≤0.0001, nonparametric Kruskal-Wallis test). N = 3 - 4 independent cultures.

**Figure 7-1: Protein and synaptic expression of Cpx-M5 mutants in mice hippocampal neuron cultures. A,** Example image (left) of a Western Blot against complexin (top) and Tubulin (bottom) for Cpx-WT rescue and Cpx mutants in Cpx-TKO hippocampal neuron cultures. Bar graph (right) corresponds to the intensity of the complexin bands in each condition normalized to WT (n = 3 independent cultures). **B,** Example images (left) and quantification (right) of immunofluorescence labeling for Complexin and V-Glut1 for continental hippocampal cultures of Cpx-TKO neurons or infected with Cpx-rescue, Cpx-M5E or Cpx-M5W. Scale bar, 10 μm. The quantification of the immunofluorescence intensity of Complexin is normalized to the immunofluorescence intensity of V-Glut1. The values are normalized to the one obtained for Cpx-WT rescue. In (B), data points represent a field of view. Data are expressed as mean ± SEM, asterisks on the graph show the significance comparisons to Cpx-rescue (* p≤0.05, **** p≤0.0001, nonparametric Kruskal-Wallis test). N = 3 independent cultures.*

