## Supplementary figures and images for "Single residues in the complexin N-terminus exhibit distinct phenotypes in synaptic vesicle fusion"

### Extended Figures

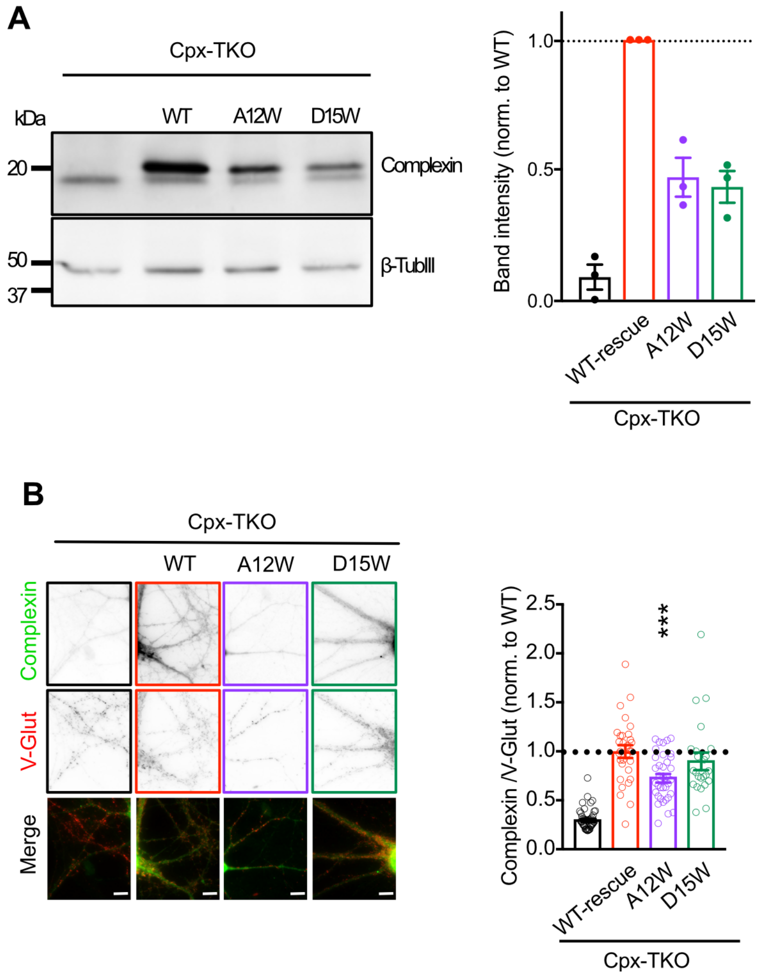

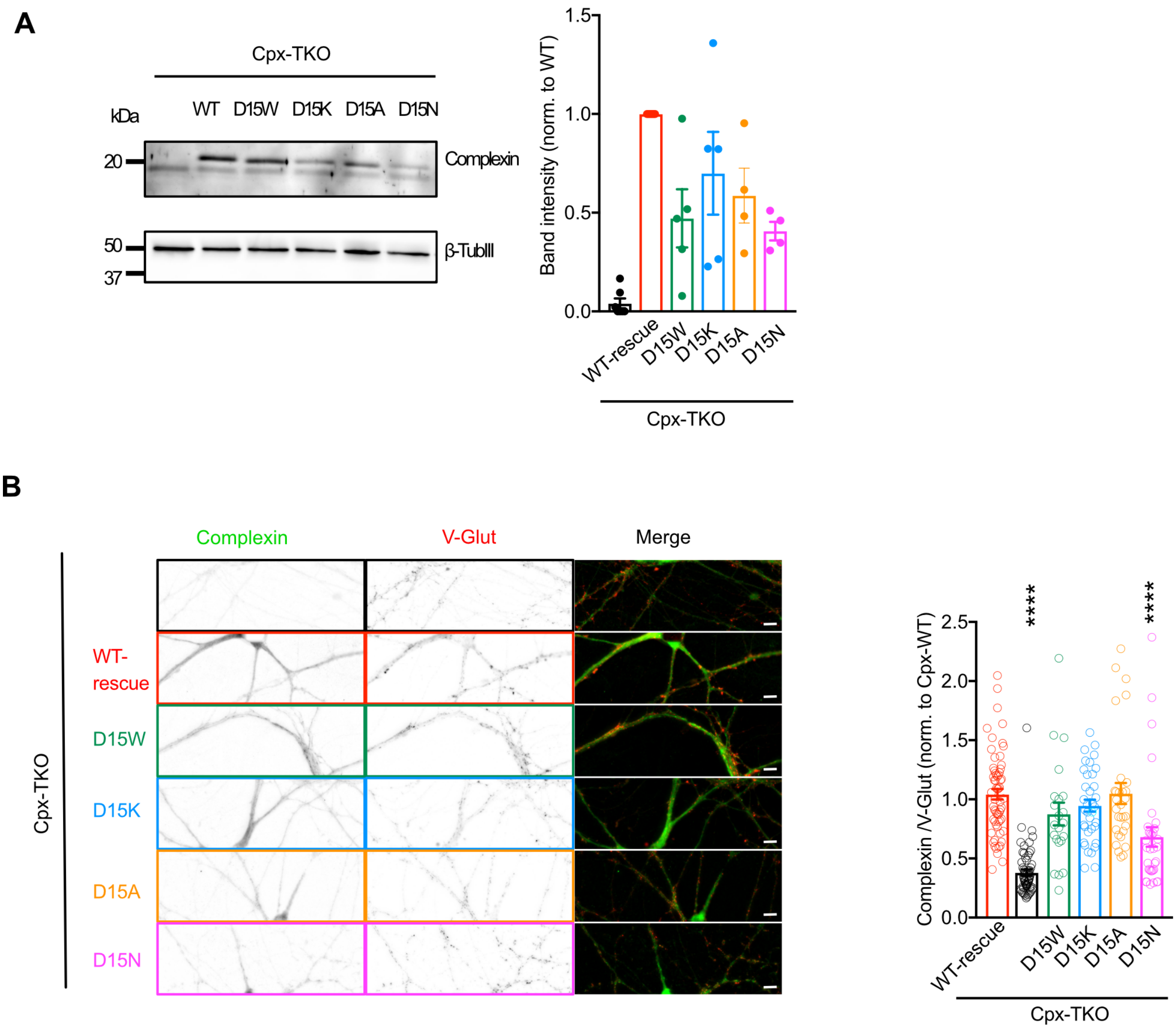

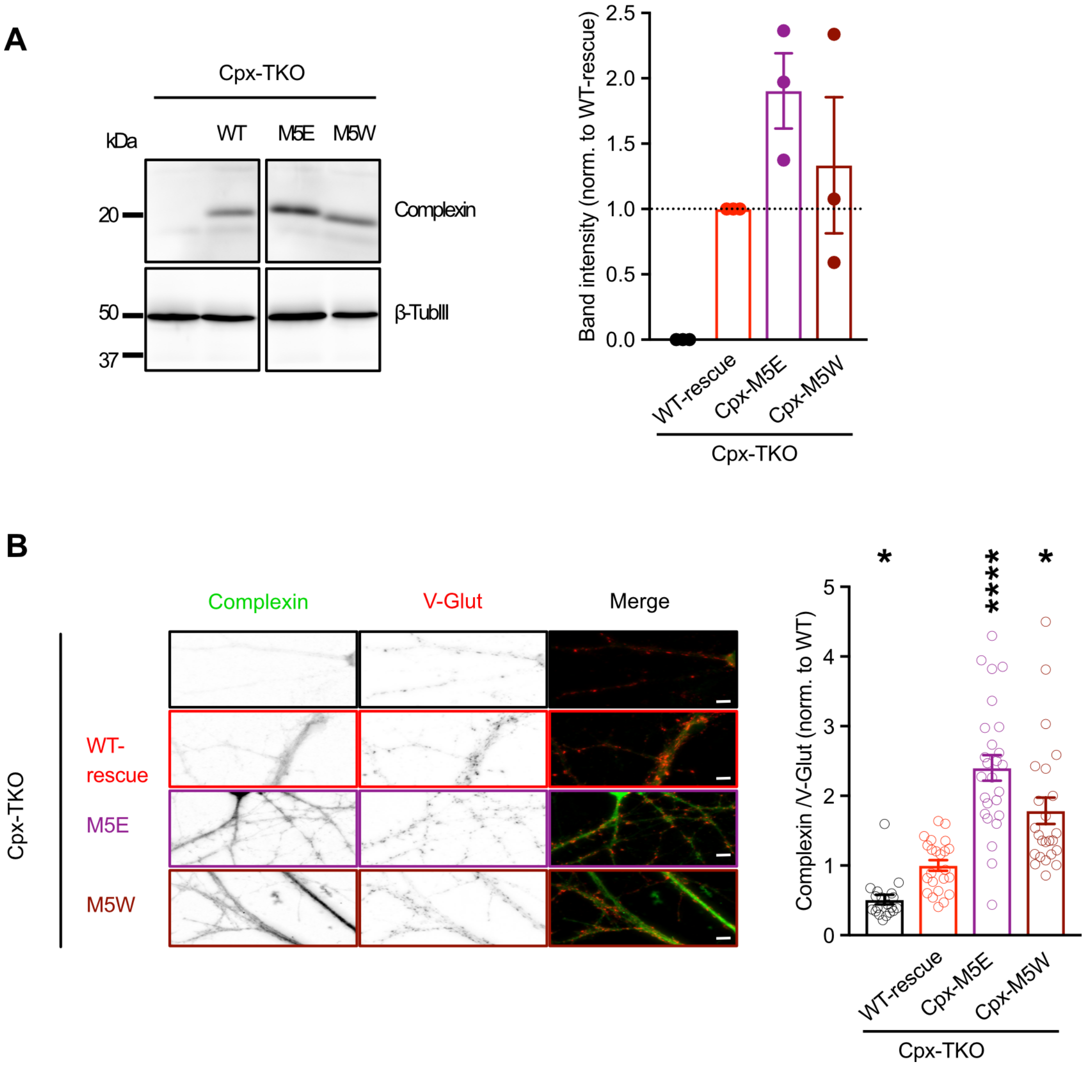
